# Emergence of Lignin-Carbohydrate Interactions During Plant Stem Maturation Visualized by Solid-State NMR

**DOI:** 10.1101/2025.01.27.635099

**Authors:** Peng Xiao, Sarah A. Pfaff, Wancheng Zhao, Debkumar Debnath, Chang-Jun Liu, Daniel J. Cosgrove, Tuo Wang

## Abstract

Lignification waterproofs and strengthens the secondary plant cell wall, while increasing the energy cost associated with releasing sugars for biofuel production. The physical association between lignin and the carbohydrate scaffold that accommodates lignin polymerization, as well as the temporally distinct roles of different lignin units and carbohydrate partners during lignification, remain largely unclear. Here we map the lignin-carbohydrate interactions by solid-state NMR in ^13^C-labeled *Arabidopsi*s inflorescence stems as secondary cell walls are formed. Analysis includes wild-type and two mutants that either selectively or globally disrupt lignin biosynthesis. Mature cell walls in the basal regions of older stems are enriched in S-lignin and carbohydrate-lignin interactions. Acetylated xylan is the dominant mediator of interactions with S-lignin, while methylated pectin unexpectedly interacts with G-lignin during early-stage lignification. The critical role of S-lignin in stabilizing carbohydrate-lignin interface is emphasized by the weak lignin-carbohydrate interactions and compromised mechanical properties of a low-S *fah1* mutant, whereas the *ref3* mutant, with low overall lignin content but a higher S/G ratio, remained unaffected. These findings demonstrate that the molecular mixing pattern, rather than lignin content, is a key determinant of the structure and properties of lignocellulosic materials.

## INTRODUCTION

Plants convert solar energy and carbon dioxide into carbohydrate-rich cell walls, collectively referred to as lignocellulose, which serves as a crucial resource for building materials, paper, textiles, biofuels, nanofibers, nanocomposites, biopolymers, and numerous other products^1–3^. Despite its functional versatility, the intricate polymer network of lignocellulose presents significant challenges for post-harvest processing and utilization in renewable energy and biomaterial applications^4,5^. Extensive efforts have focused on engineering crops with improved biopolymer composition and structure, as well as optimizing enzymatic and chemical methods for digestion and treatment of these biomaterials^6–8^. However, a critical barrier remains: our limited understanding of cell wall architecture at the molecular level, particularly the polymorphic structures and physical interactions of complex carbohydrates with other biomolecules in their native cellular context. Notably, little is known about early stages of secondary wall formation in maturing cells.

The plant secondary cell wall constitutes the majority of lignocellulosic biomass and is crucial for providing structural integrity, strength, and rigidity to specialized plant cells such as xylem vessels, fibers, and sclereids with thick lignified cell walls^9^. Its formation is a highly orchestrated process that relies on the regulation of genes encoding enzymes and transcription factors^10–12^. This process commences following the completion of primary cell wall development, involving the deposition of cellulose microfibrils, hemicelluloses, and lignin to the thickening cell wall. In many secondary walls, cellulose microfibrils are deposited sequentially into three distinct layers (S1, S2, and S3), which differ in microfibril orientation and organization^13,14^. Microfibrils commonly aggregate in secondary walls to form larger fibrils, or macrofibrils, measuring tens of nanometers across^15,16^. The assembly of macrofibrils also involves hemicelluloses, such as xylan and glucomannan, which may modulate microfibril aggregation and facilitate the attachment of lignin to the carbohydrate scaffold^16,17^.

As a hallmark of secondary cell wall maturation, lignification involves the polymerization of monolignols—phenolic compounds synthesized via the phenylpropanoid pathway and likely transported into cell walls through diffusion^18,19^. These monolignols undergo oxidative coupling catalyzed by enzymes such as peroxidases and laccases, which may deplete monolignols in the cell wall, thereby sustaining the concentration gradient necessary for continuous monolignol transport^19,20^. Once incorporated into the carbohydrate matrix, lignin enhances the mechanical strength and hydrophobicity of secondary cell walls, as well as their resistance to mechanical stress, pests, and pathogens^21,22^. However, the recalcitrant nature of the lignin polymer poses challenges for biomass processing, particularly in applications such as pulping and biofuel production^5,6^. Meanwhile, considerable effort has been put towards effectively converting residual lignin into valuable products, but lignin valorization has been hindered by the structural heterogeneity and unfavorable physical and chemical properties of these polymers^4,23^.

The challenge of understanding the molecular architecture of mature secondary cell walls has been partially addressed through the application of solid-state NMR spectroscopy across diverse plant species, including poplar, eucalyptus, *Arabidopsis*, spruce, maize, switchgrass, rice, *Brachypodium*, and sorghum^24^. This approach has led to five advances in our understanding of the structural principles governing the interactions of carbohydrates and lignin in plant secondary cell walls. The chemical and conformational structure of xylan determines its binding specificity within the cell wall, with an evenly substituted xylan pattern and a flat-ribbon conformation, termed two-fold xylan, being essential for deposition onto the cellulose surface^25,26^. When this flat-ribbon structure is disrupted, such as by irregular substitutions observed in sorghum, non-flat (three-fold) xylan can still serve as binding modules for disordered regions of cellulose microfibrils^27^. Furthermore, three-fold xylan predominantly acts as an interface between cellulose fibrils and lignin nanodomains^28–30^. The physical packing of the lignin-xylan interface is primarily stabilized by electrostatic interactions involving polar motifs, with syringyl (S) lignin playing a more prominent role than guaiacyl (G)^30^. Additionally, while xylan, particularly the three-fold conformation, is the primary interactor with lignin, cellulose also participates in these interactions in mature stems, where molecular crowding promotes contact between the components^28,31,32^.

Given the dynamic nature of the lignification process, the next key question to address is how the nanostructure of the lignin-carbohydrate interface evolves during the formation of secondary cell walls. Here we choose maturing *Arabidopsis* inflorescence stems as the model system, and characterize different segments of the stems grown to varying ages and heights, capturing the progression of secondary cell wall formation under standardized growth conditions. Wild-type (WT) stems were compared with two mutants, *fah1-2* and *ref3-3*, which have altered lignin content and composition. The *ref3-3* mutant, carrying a mutation in cinnamate 4-hydroxylase (C4H)^33^, exhibits reduced overall lignin content but a higher S/G ratio^34–36^. The *fah1-2* mutant has an undetectable transcript level of ferulate 5-hydroxylase (F5H), which hydroxylates coniferyl alcohol/coniferaldehyde (the G-lignin precursor) at its C-5 position to form S-monolignols, leading to defective S-lignin biosynthesis but elevated or unchanged G subunit content^37–40^. By combining solid-state NMR spectroscopy with ^13^C enrichment and genetic mutants, we managed to resolve the temporal functions of different lignin types during the formation of secondary cell walls, and the different carbohydrates that serve as binding partners.

## RESULTS

### Aged inflorescence stems exhibit higher S-lignin content and carbohydrate interactions

To assess whether lignification depends on the age of the inflorescence stem, we grew three sets of uniformly ^13^C-labeled wild-type (WT) *Arabidopsis* stems to different heights and harvested 2-cm basal segments for comparison (**Fig. 1a**). The 1D ^13^C cross-polarization (CP) spectra, which selectively detect the rigid fraction of the cell walls, showed a sequential increase in the lignin signal from the basal samples of WT08A, WT12A, and WT16A (**Fig. 1b**), revealing that lignin content increased with the age and developmental stage of the inflorescence stems. Indeed, the increase in lignin content was consistently maintained even as the stem grew to a height of 32 cm (**Supplementary Fig. 1**). Intensity analysis, comparing the integral of the aromatic region to the whole spectrum, demonstrated that lignin content increased from 3% to 5% of the rigid fraction between WT08A and WT16A (**Fig. 1c**). The crystalline cellulose content in WT08A was also low, as indicated by reduced intensities at 89 ppm (carbon 4) and 65 ppm (carbon 6). Spectral deconvolution revealed an increase in the average content of interior chains within cellulose microfibrils, rising from 27% in WT08A to 30% in WT12A and WT16A, indicating enhanced cellulose crystallinity as the plant reached a height of 12 cm (**Supplementary Fig. 2**).

**Figure 1.**
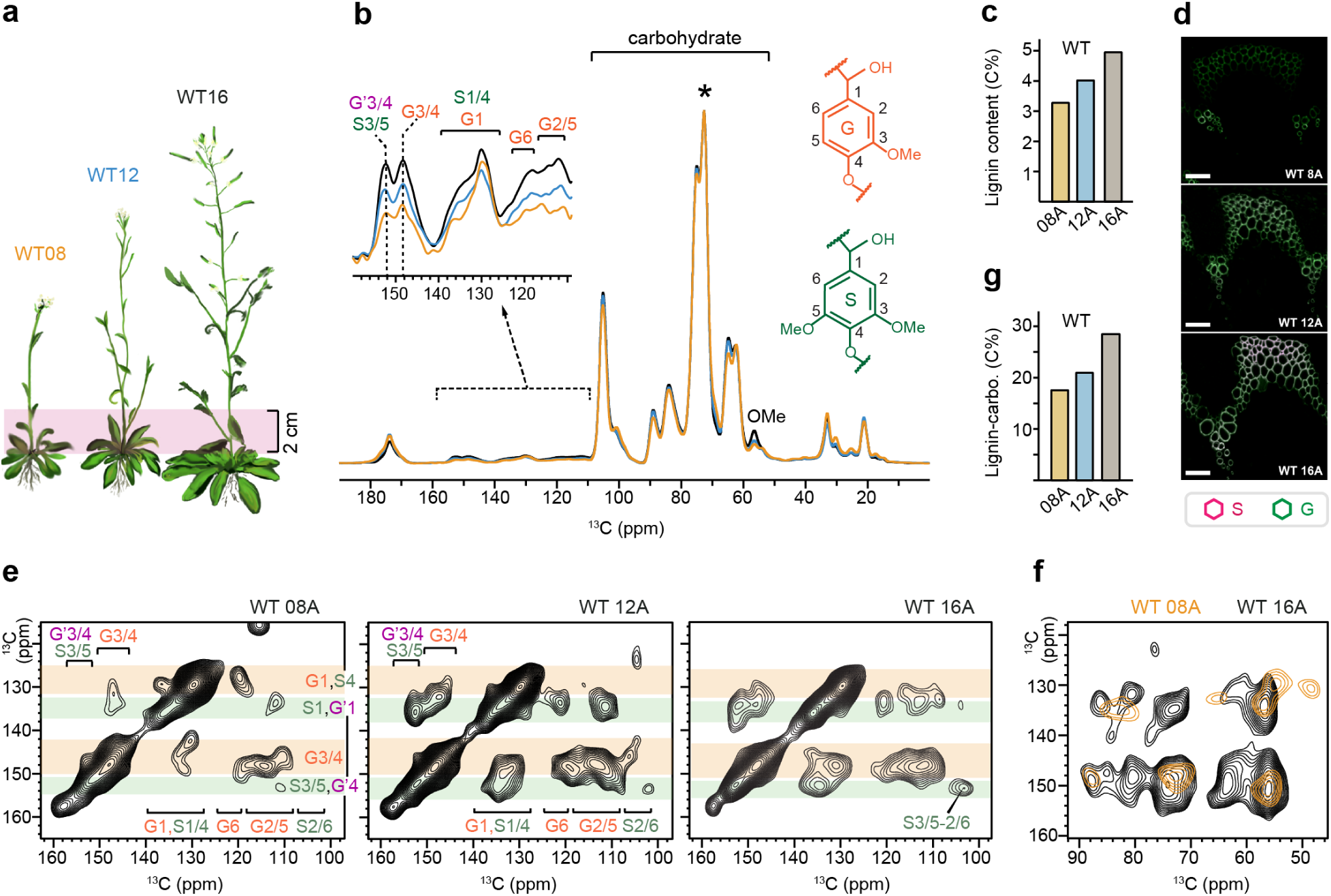
Lignin composition and carbohydrate interactions vary in wild-type inflorescence stems. (**a**) Three samples collected from the basal 2-cm segment of wild-type (WT) of *Arabidopsis* grown to different heights. (**b**) 1D ^13^C CP spectra collected on the basal segment of *Arabidopsis* grown to 8 cm (WT08A, yellow), 12 cm (WT12A; cyan) and 16 cm (WT16A; black) height. All spectra were normalized by the 72 ppm peak (asterisk), the highest signal of the carbohydrate region. Zoomed-in view was provided for the lignin aromatic region showing the changes in the overall lignin content. S: syringyl unit. G and G’: guaiacyl units. For example, G1 represents the carbon 1 of G unit. (**c**) The overall lignin content in the rigid fraction estimated from the 1D CP spectra. (**d**) Mäule staining shows progression of lignification in SCW-forming tissues during inflorescence maturation. Merged fluorescence of two-channel images shows G-lignin (green) and S-lignin (pink). Scale bars = 50 µm. (**e**) 2D ^13^C-^13^C correlation spectra measured on the three samples with a short DARR mixing time of 0.1 s, showing intramolecular short-range cross-peaks. Key regions of S and G carbons are marked on the horizontal dimension (ω_2_ dimension). The vertical dimension (ω_1_) was highlighted for G-dominant bands (orange) and S-dominant bands (green). (**f**) Overlay of the lignin-carbohydrate interaction region of two 2D ^13^C-^13^C correlation spectra measured on WT08A and WT16A samples with a long PDSD mixing time of 1.0 s for detecting intermolecular long-range cross-peaks. (**g**) Relative abundance of lignin-contacted carbohydrates estimated from the peak volumes of intermolecular cross-peaks in panel (f).

As noted in **Fig. 1b**, the abundance of lignin methoxy groups (OMe) was substantially lower in WT08A and WT12A compared to WT16A, indicating that not only lignin content but also the syringyl-to-guaiacyl (S/G) ratio increased in the older 16-cm stem. Intensity analysis suggested that the S content increased from 25–26% in WT08A and WT12A to 33% in WT16A (**Supplementary Fig. 3a, b**). This was further confirmed using both microscopic and spectroscopic methods. First, Mäule staining of stem cross-sections enabled the visualization of G and S lignin distribution based on their distinct fluorescence emission profiles (**Fig. 1d** and **Supplementary Fig. 4**). In WT08A, lignified xylem walls were observed, primarily composed of G-lignin, while WT12A exhibited increased lignification in both xylem and interfascicular fiber cells, with a higher proportion of S-lignin becoming apparent. In WT16A, the overall lignin content was greater across both cell types, with more pronounced tissue-specific lignification patterns, characterized by G-lignin predominating in vessel cells and S-lignin in fiber cells. Second, 2D ^13^C-^13^C correlation spectra showed that the characteristic S3/5-2/6 cross-peak at (153, 104 ppm) was prominent in WT16A but much weaker in WT12A and almost absent in WT08A (**Fig. 1e**). In contrast, the G-rich spectral bands dominated the WT08A spectrum. These results revealed the predominant deposition of G units into the cell wall in the earlier stages, while more S units, with a higher level of methoxy substitution, were incorporated as the stem matured.

Another notable observation is that lignin in the WT16A sample exhibited significantly more extensive interactions with cell wall carbohydrates. This is demonstrated by the stronger and more abundant intermolecular cross peaks between lignin aromatic carbons (120-160 ppm) and distinct carbohydrate carbons (60-90 ppm), as shown in **Fig. 1f**. Intensity analysis, presented in **Supplementary Fig. 5**, revealed that carbohydrates in close proximity to lignin accounted for only 18% of all carbons within approximately 1 nm of lignin aromatics in the WT08A sample, but this percentage increased to 29% in the WT16A sample (**Fig. 1g**). Thus, as the inflorescence stem developed, more S residues were incorporated, leading to an increase in lignin methoxy substitutions and tighter physical packing of lignin with polysaccharides in the cell wall. This observation supports the role of lignin’s methoxy groups in stabilizing associations with carbohydrates, primarily through interactions with xylan and, to a lesser extent, with cellulose, as recently shown^29–31^.

### The basal region of the inflorescence stem exhibits increased lignification

To assess lignification along the same inflorescence stem, we analyzed three 2-cm segments from the basal region of a WT plant grown to a height of 16 cm (basal A, B, C; **Fig. 2a**). Compared to the lower two segments, aromatic peak intensity decreased in segment C (WT16C), located 4-6 cm from the base (**Fig. 2b**). Closer to the apex, lignin signals became increasingly faint due to the lower abundance of secondary cell walls in younger tissues (**Supplementary Fig. 6**). Segments A and B showed similar lignin profiles, with segment A displaying a slight increase in the 153-ppm peak, dominated by S3/5 carbons of S-units, with minimal contribution from a G-residue variant (G’). Meanwhile, the G3/4 peak at 148 ppm remained unchanged (**Fig. 2b**). Thus, the lowest (most mature) segment showed a moderate increase in the S/G ratio of lignin composition, with S-lignin comprising 31%, a slightly higher percentage than the 29% observed in the two upper segments (**Supplementary Fig. 3c, d**).

**Figure 2.**
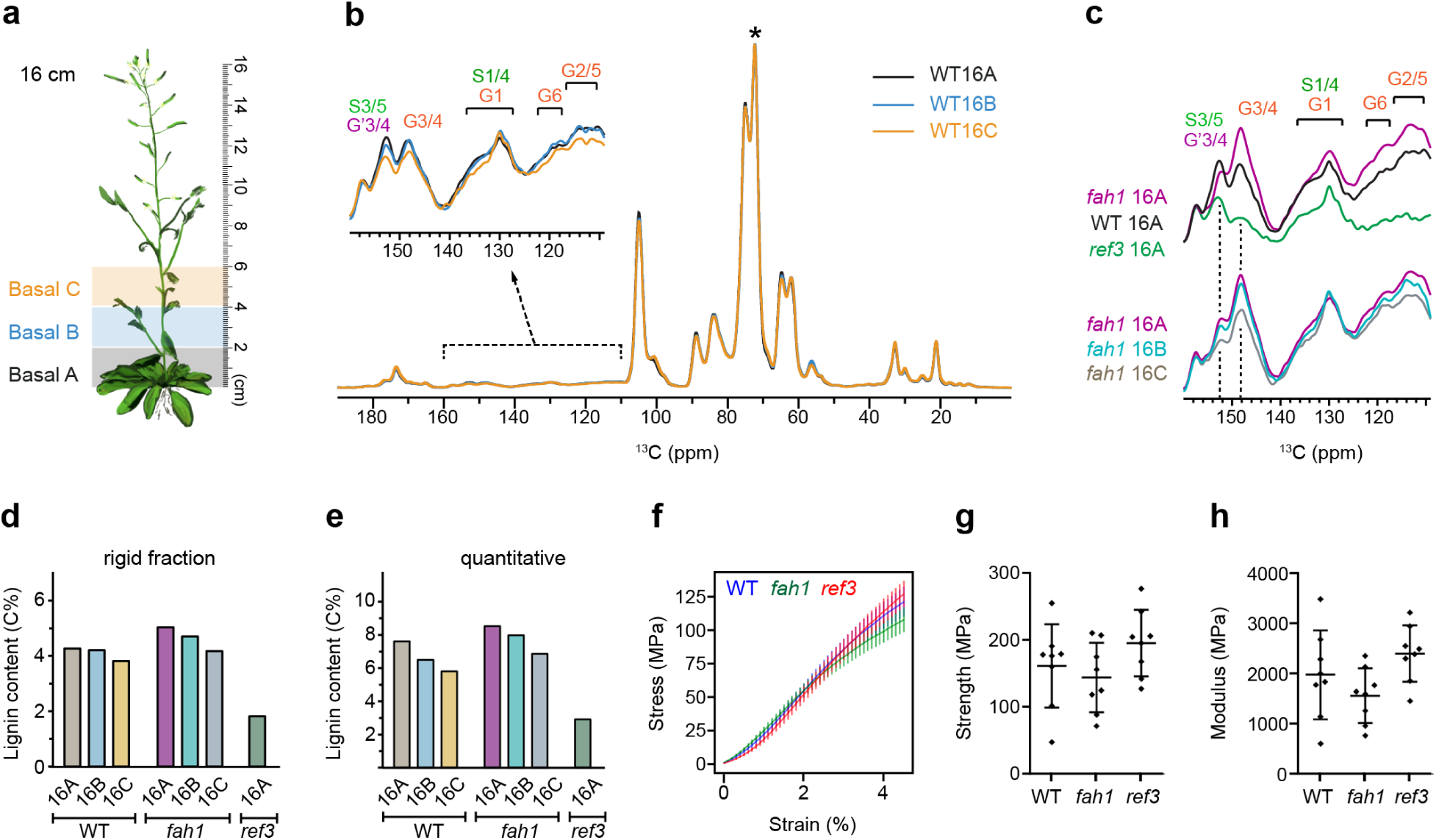
Lignification and tensile mechanical properties of mutant inflorescence stems. (**a**) Illustration of the positions of basal segments A (0-2 cm), B (2-4 cm), and C (4-6 cm) cut from the *Arabidopsis* plant. (**b**) 1D ^13^C CP spectra collected on the three segments of the same stem of WT *Arabidopsis* grown to 16 cm. The samples are referred to as WT16A (black), WT16B (blue), and WT16C (yellow). All spectra are normalized with respect to the highest 72-ppm carbohydrate peak (asterisk). Zoomed-in view was provided for the lignin aromatic region showing the changes in the rigid lignin content. (**c**) Comparison of the aromatic region of 1D ^13^C CP spectra collected on segment-A from the 16-cm plants of WT *Arabidopsis* and two mutants *fah1* and *ref3*. Overlap of the spectra of the A, B, and C segments of *fah1* is also shown in the bottom panel. All spectra are normalized by the highest 72-ppm carbohydrate peak although the carbohydrate region is not included in this figure. (**d**) Lignin content in the rigid fraction detected by CP within different WT and mutant samples. (**e**) Quantitative analysis of overall lignin content from ^13^C DP spectra measured with long recycle delays of 35 s. (**f**) Stress-strain curves of monotonic tensile loading tests indicate similar mechanical properties for all three analyzed genotypes of basal inflorescence stems. Error bars represent s.d. (**g**) Stem segment breaking strength and (**h**) modulus at the 4.5%-5% strain interval for the basal inflorescence stems of different genotypes. Both scatterplots show the mean with a horizontal line while the error bars represent s.d. (n = 8).

The increase in lignin content along the apical-to-basal axis was more clearly observed in the *fah1-2* mutant with a defective ferulate 5-hydroxylase (F5H) gene^37,38,41^, where lignin content progressively decreased in segments A, B, and C (**Fig. 2c** and **Supplementary Fig. 7**). This mutant is known to deplete most S units from lignin in *Arabidopsis*, resulting in the G3/4 peak dominating the spectra^41^. In contrast, the *ref3-3* mutant carrying a mutation in C4H^34–36^, which exhibits reduced overall lignin content but an increased S/G ratio, showed spectra dominated by the S3/5 peak (**Fig. 2c**). Intensity analysis of CP spectra indicated that lignin content sequentially decreased from 4.4% to 4.2% and then to 3.8% in the rigid fraction of segments A, B, and C of the WT sample, respectively (**Fig. 2d**). A similar sequential decrease (5.0%, 4.7%, and 4.2%) was observed for the *fah1-2* mutant, while the *ref3-3* mutant exhibited a substantially lower lignin content of only 1.8%. Since lignin contains an appreciable amount of both mobile and rigid fractions, whereas polysaccharides are primarily found in the rigid fraction, CP spectra, which selectively detect rigid molecules, may underestimate the total lignin content. To obtain a more accurate measure, we performed a detailed analysis using quantitative ^13^C direct polarization (DP) spectra with long recycle delays of 35 seconds to ensure unbiased detection of all carbon atoms in the sample (**Supplementary Fig. 8**). The overall lignin content was nearly twice as high as the value reported by CP data, and both the WT and *fah1-2* samples exhibited a more pronounced trend of decreasing lignification content when moving away from the base of the stem (**Fig. 2e**).

Tensile loading tests of basal inflorescence segments, presented as stress-strain curves normalized to the dry weight of the stretched cell wall material, revealed that the *fah1* mutant was slightly weaker, while WT and *ref3* exhibited nearly identical mechanical behavior (**Fig. 2f**). No statistically significant differences were found between the three genotypes with respect to tensile strength (**Fig. 2g**) and modulus within the 4.5-5.0% strain interval (**Fig. 2h**). These results are notable given that *fah1* exhibited a slightly higher lignin content, predominantly G-lignin, relative to WT, whereas *ref3* contained less than half the lignin content, which was primarily S-lignin (**Fig. 2c, d**). These findings indicate that variations in the total lignin content, or the specific contribution of G-lignin, did not substantially influence the tensile mechanical properties of inflorescence stems. Alternatively, it is plausible that S-lignin plays a more significant role than G-lignin in mechanical stabilization. This could explain why the low-S *fah1* mutant is slightly weaker (at 4% strain), while the *ref3* mutant, despite its considerably lower overall lignin content, maintains tensile mechanics comparable to the WT.

### Mapping the lignin composition and carbohydrate contacts in WT and mutant stems

When projected to 2D refocused J-INADEQUATE experiments for better spectral resolution, both WT16A and *ref3-*16A samples exhibited a mixture of syringyl (S) units and their Cα-oxidized forms (S’), along with guaiacyl (G) residues, whereas the *fah1*-16A sample exclusively displayed peaks corresponding to G units (**Fig. 3a**). This distinction was traced through the characteristic carbon signals at the C3 and C4 positions. For S units, the single-quantum (SQ) chemical shifts were observed at 154 ppm (S3/5) and 134 ppm (S4), with a corresponding double-quantum (DQ) shift at 288 ppm, which represents the sum of the two SQ shifts from directly bonded carbons. The SQ chemical shifts for G units ranged between 142-154 ppm (G3/4), with DQ shifts of 290-300 ppm. We summarized the available chemical shift data from recent studies (**Fig. 3b**) to aid in resonance assignments and will serve as a valuable reference for future research.

**Figure 3.**
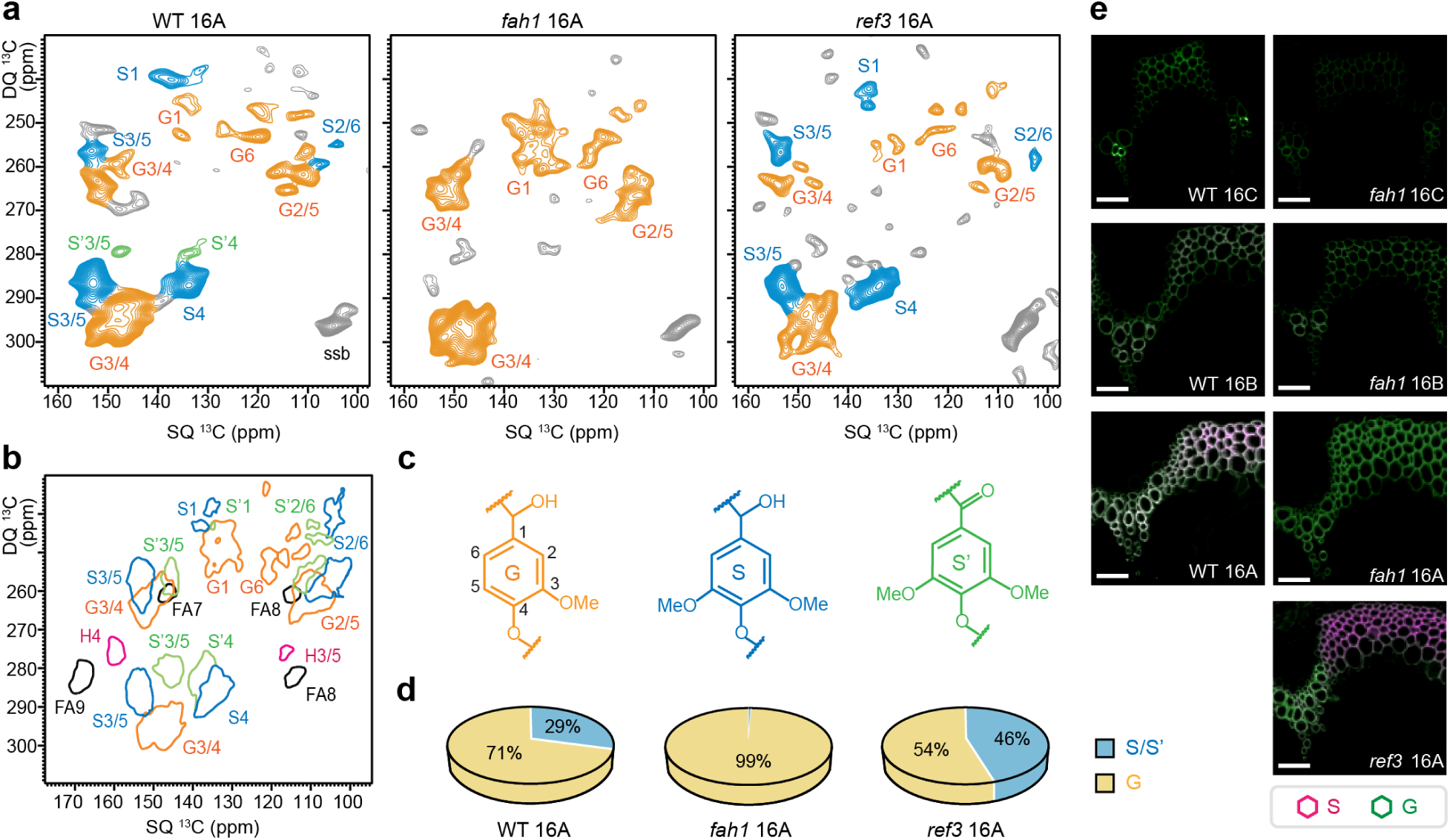
Monolignol composition in wild-type *Arabidopsis* and lignin mutants. (**a**) A comparison of lignin regions from 2D ^13^C CP refocused J-INADEQUATE spectra of WT16A (left), *fah1-*16A (middle), and *ref3-* 16A (right). The signals are color-coded to represent syringyl (S, blue), guaiacyl (G, orange), and oxidized syringyl (S’, green) units. (**b**) A simulated distribution map of resolvable NMR signals for different lignin monomer units, generated using data from multiple plant species including eucalyptus, poplar, spruce, and maize. H: *p*-hydroxyphenyl unit, FA: ferulate. (**c**) Representative chemical structures of the G, S, and S’ units. (**d**) Molar fractions of G (yellow) and combined S/S’ (blue) units in the three samples. Estimations were based on the peak volumes corresponding to carbon 3/5 and 4 of the S/S’ units, and carbon 3/4 signals of the G units. (**e**) Representative images of Mäule-stained cross-sections of *Arabidopsis* stems. The merged fluorescence images show the distribution of G lignin (green) and S lignin (pink) in lignified tissues of the basal segments of the inflorescence stems. Scale bars represent 50 μm.

Analysis of the resolved peak volumes indicated that the combined fraction of S and S’ units (**Fig. 3c**) accounted for 29% of the lignin signals in WT16A, with G units making up the remaining 71%, resulting in an S/G ratio of 0.41 (**Fig. 3d**). A 4% discrepancy with the value obtained from 1D spectra indicates the error margin introduced by utilizing different spectral methods in lignin content analysis. The S/G ratio decreased to 0.25 in the basal-A segment of stems that had grown to heights of 19 cm and 32 cm (**Supplementary Fig. 9**), which is consistent with the range of 0.22-0.35 reported for mature *Arabidopsis* plants^35–38,42^. Although the *ref3-3* mutant had a reduced overall lignin content, it exhibited an almost doubled S/G ratio (0.85 in *ref3-*16A) compared to the wild-type sample. Consistently, Mäule staining of the *fah1* mutant revealed exclusive deposition of G-lignin, with no detectable S monomers across all SCW-forming cell types (**Fig. 3e** and **Supplementary Fig. 4**). A clear gradient of lignin deposition was observed along the basal region of the stem, progressing from the 16C segment to the 16A segment. In the *ref3* mutant, the most distinct cell-specific separation of lignin monomers was evident, with G-lignin predominantly localized in vessel cells and S-lignin more prominent in fiber cells.

### Evolution of lignin-carbohydrate packing in different segments of WT and mutant stems

The physical packing interactions occurring between lignin and polysaccharides were probed through a series of dipolar-gated 2D ^13^C-^13^C correlation experiments (**Fig. 4a**, **Supplementary Figs. 10-12**). A long mixing period (1.0 s) facilitated further polarization transfer, uncovering numerous long-range intermolecular cross peaks that represent packing interactions on the subnanometer scale, which were not observed in the short-mixing (0.1 s) spectra (**Fig. 4a**). *ref3-* 16A exhibited a different spectral pattern compared to WT16A and *fah1-*16A: *ref3-*16A showed fewer intramolecular cross peaks in the 0.1 s spectrum, likely due to its compromised lignin synthesis, but displayed strong long-range correlations in the 1.0 s spectrum. This observation indicates that, despite the lower lignin quantity, the remaining lignin was well-packed with carbohydrates in the *ref3* mutant (**Supplementary Fig. 13**).

**Figure 4.**
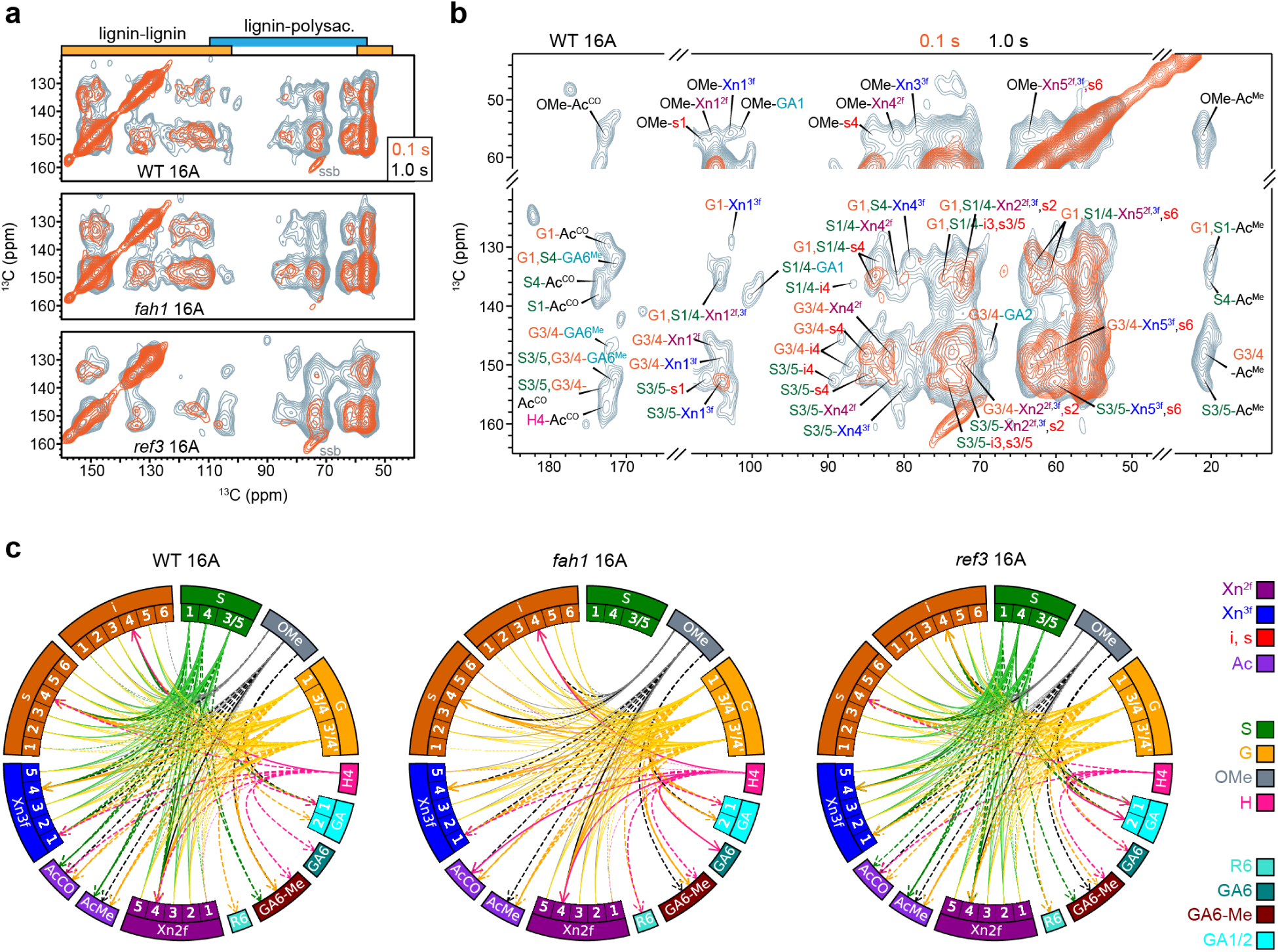
2D correlation spectra showing intermolecular interactions. (**a**) Overlap of 2D dipolar-gated ^13^C-^13^C correlation spectra measured with a short mixing time (0.1 s, orange) and a long mixing time (1.0 s, grey). The latter showed additional signals coming from long-range intermolecular interactions. The basal-A segments of WT and two lignin mutants are included. Representative spectral regions of lignin-lignin and lignin-polysaccharide cross peaks are marked. (**b**) Key spectral regions of the WT16A sample showing lignin-carbohydrate interactions. Xn^3f^: 3-fold xylan; Xn^2f^: 2-fold xylan; Ac: acetyl; OMe: lignin methoxy; i: interior cellulose; s: surface cellulose; GA: galacturonic acid; R: rhamnose; CO: carbonyl. For example, the label S1/4-s4 represents the cross peak observed between the carbon 1 and 4 sites of S-lignin with the carbon 4 of surface cellulose. (**c**) Chord diagrams visualizing intermolecular lignin-carbohydrate interactions observed in WT16A, *fah1-*16A and *ref3-*16A. Solid lines and dashed lines represent cross-peaks identified in the short-range and long-range spectra, respectively. Thick lines with arrows indicate unambiguous, site-specific cross-links and narrow lines without arrow indicate links from convoluted cross-peaks.

The observed interactions included those between lignin methoxy (OMe) groups and various carbohydrate carbons, such as those found in 2- and 3-fold xylan, surface cellulose chains, acetyl groups, including both their methyl and carbonyl carbons, and even some pectic polymers. Key aromatic carbons of S and G units also showed extensive cross peaks with these carbohydrate carbons. S and G units both interacted with the surface chains of cellulose extensively but exhibited distinct patterns of interactions with internal chains. Interactions between S/G units and the interior chains of cellulose were also detected (**Fig. 4b**), as evidenced by strong cross-peaks such as S3/5-i4 (152, 89 ppm) and G3/4-i4 (145, 89 ppm). These cross-peaks reflect the tight physical packing within the surface-proximal interior layer of cellulose microfibrils, with components colocalized within a nanometer scale. A relatively weak but unambiguous cross-peak of S1/4-i4 (136, 87 ppm) was also observed, likely arising from relayed polarization transfer from S-lignin to either deeply embedded, inaccessible interior cellulose chains or interior chains with structural alterations due to their interactions with hemicellulose^43,44^.

A contact-map representation of a total of 921 lignin-carbohydrate interactions observed in seven samples revealed distinct differences in the spatial organization of these polymers between wild-type and mutant stems (**Fig. 4c**; **Supplementary Fig. 14**). Each sample showed approximately 60 S-carbohydrate interactions, except in the S-depleted *fah1* samples, 60-70 G-carbohydrate interactions, 20-30 OMe-carbohydrate cross peaks, and a few H-carbohydrate cross peaks (**Supplementary Table 1**). In WT16A, S-lignin primarily interacted with xylan (Xn^3f^, Xn^2f^, AcCO, and AcMe) and cellulose (i and s), with only four cross peaks observed for carbon 6 of methylated galacturonic acid (GalA) in pectin (GA6-Me) and carbon 1 of GalA (GA1). In contrast, G-units and the weakly resolved carbon 4 of H-lignin (156-161 ppm) showed broader correlations with pectin, including methylated and unmethylated GalA units and Rha, spanning homogalacturonan (HG) and rhamnogalacturonan I (RG-I). In *fah1-*16A, depletion of S-units significantly reduced the number of lignin-carbohydrate cross peaks (**Fig. 4c**), which might explain the weaker wall of the *fah1* mutant (**Fig. 2f, h**). For instance, carbon 1 of surface and interior cellulose chains exhibited only a single weak interaction with G-lignin in *fah1-*16A, compared to four interactions with both S and G units in WT16A. The polymer contact map in *ref3-*16A was nearly restored (**Fig. 4c**), resulting in segment strength and modulus comparable to the WT sample (**Fig. 2g, h**).

To estimate the percentage of lignin and carbohydrates in contact with each other, we introduced a straightforward approach by comparing the integrals of key spectral regions (**Fig. 5**; **Supplementary Fig. 15**). The heatmap intensities represent the percentage of carbohydrate carbons found within a 1 nm distance from reference lignin sites. For instance, **Figure 5a** shows four analyses of these interactions: (1) all lignin carbons (S/G units and lignin methoxy groups) interacting with carbohydrate carbons (including acetyls), (2) all lignin carbons interacting with carbohydrate carbons, excluding acetyls, (3) lignin ring carbons (S/G units) interacting with carbohydrate carbons, excluding acetyls, and (4) lignin methoxyl groups interacting with carbohydrate carbons, excluding acetyls. Additional detailed analyses were performed as needed to further map the interactions between various lignin units and structural motifs with the acetyl groups (**Fig. 5b**) and ring carbons (**Fig. 5c**) in carbohydrates.

**Figure 5.**
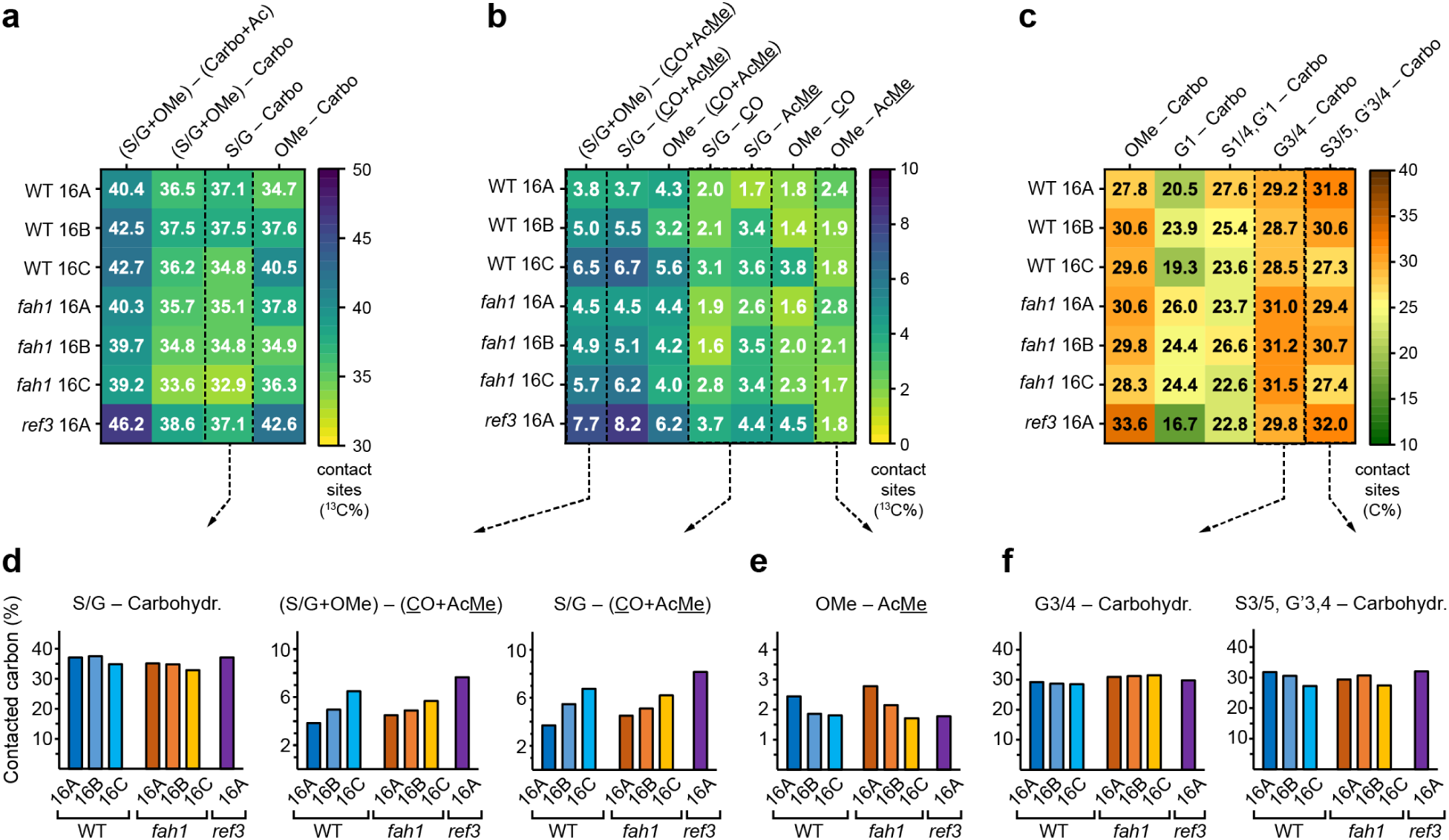
Heatmap presentation of lignin-carbohydrate interactions in seven *Arabidopsis* samples. (**a**) Overall lignin-carbohydrate interactions analyzed in four different ways depending on whether the lignin methoxy and carbohydrate acetyls are included or not. S/G: lignin aromatic carbon only; OMe: lignin methoxy; carbo: carbohydrate carbons with acetyls excluded; Ac: acetyl. The heatmap intensity represents the contacted carbon percentage calculated from cross peak integrals extracted from 2D dipolar-gated ^13^C-^13^C correlation spectra with long mixing time (1.0 s). (**b**) Analysis of lignin-acetyl interactions. (**c**) Interactions between resolvable lignin sites and carbohydrate carbons with acetyls excluded. Key lignin-carbohydrate interactions that evolve in different inflorescence segments are shown separated in panels (**d**-**f**).

Among the seven samples analyzed, *ref3-*16A exhibited the strongest lignin-carbohydrate interactions, with 46.2% of all carbons surrounding lignin within a 1 nm distance being occupied by carbohydrate carbons (**Fig. 5a**). Interestingly, despite the relatively low level of lignification in *ref3-*16A (**Fig. 2c**), this may have reduced lignin self-aggregation, allowing for a greater proximity of carbohydrate polymers. Additionally, the elevated syringyl/guaiacyl (S/G) ratio and high methoxy group content in *ref3-*16A lignin may have further contributed to this enhanced interaction. Similarly, in the WT stems, the C segment contained the lowest lignin content (**Fig. 2d, e**) and exhibited better integration of lignin with carbohydrates (**Fig. 5a**).

The opposite trend was observed when either the acetyl groups of carbohydrates or the methoxy groups of lignin were excluded from the analysis (second and third columns of **Fig. 5a**). This indicates that these motifs behaved differently from the bulk carbons of lignin and carbohydrates, serving as the primary drivers of distinguishing intermolecular packing interactions. This was confirmed by the high level of methoxy-carbohydrate contact, 40.5%, in WT16C (**Fig. 5a**). Additionally, the extent of lignin-acetyl contact increased progressively across segments A, B, and C in both WT16 and *fah1*-16 stems (**Fig. 5d** and **Supplementary Fig. 15**). In contrast, direct methoxy-acetyl contacts decreased sequentially across the three segments (**Fig. 5e**), aligning with the increasing methoxy content in the lignin structure in the more basal stem segments.

In both WT and *fah1* samples, interactions between G3/4 units and carbohydrates remained constant across the three basal segments (**Fig. 5f**). However, interactions between S units and carbohydrates decreased in the higher segments. This trend is evident in the data for S1/4 and G’1-carbohydrate, as well as S3/5 and G’3/4-carbohydrate interactions, which sequentially diminished from segments A to C (**Fig. 5c, f**), though minor contributions from G units are also present in these measurements. In comparison to WT16A, the S units in the higher segment WT16C are not only less abundant but are also, on average, spatially further from carbohydrates.

### Identification of pectin-lignin interactions in inflorescence stems

We distinguished the carbonyl (CO) signals originating from the acetyl groups (-OCOMe) of either acylated xylan or pectin, e.g., galacturonic acid (GalA) units, at 173.5 ppm, the carboxylates (-COO⁻ or -COOH) from unmethylated GalA at 176 ppm, and the methyl esters (-COOMe) from methylated GalA at 172-168 ppm (**Fig. 6a**). This enabled us to assess the contributions of various carbohydrates to lignin interactions by tracking CO chemical shifts (**Fig. 6b**). Most of the long-range interactions with lignin occurred via acetyl groups, predominantly found in xylan in these samples. Additional interactions were observed at the carboxylate and methyl ester sites of pectin. Notably, interactions between pectin methyl esters and G1 and G6 were detected in WT16B and WT16C but not in WT16A. This indicates strong interactions between methylated pectin and G-lignin in younger segments of the WT inflorescence stem, where the lignin is rich in G units. In contrast, unmethylated pectin exhibited minimal interaction with lignin, with only weak cross-peaks observed between carboxylates and G units, primarily in WT16A, *fah1-*16A, and *ref3-*16A, which correspond to older stem segments. This may reflect the temporal progression of pectin demethylation, occurring after the deposition of methylated pectin polymers in the cell wall^45,46^.

**Figure 6.**
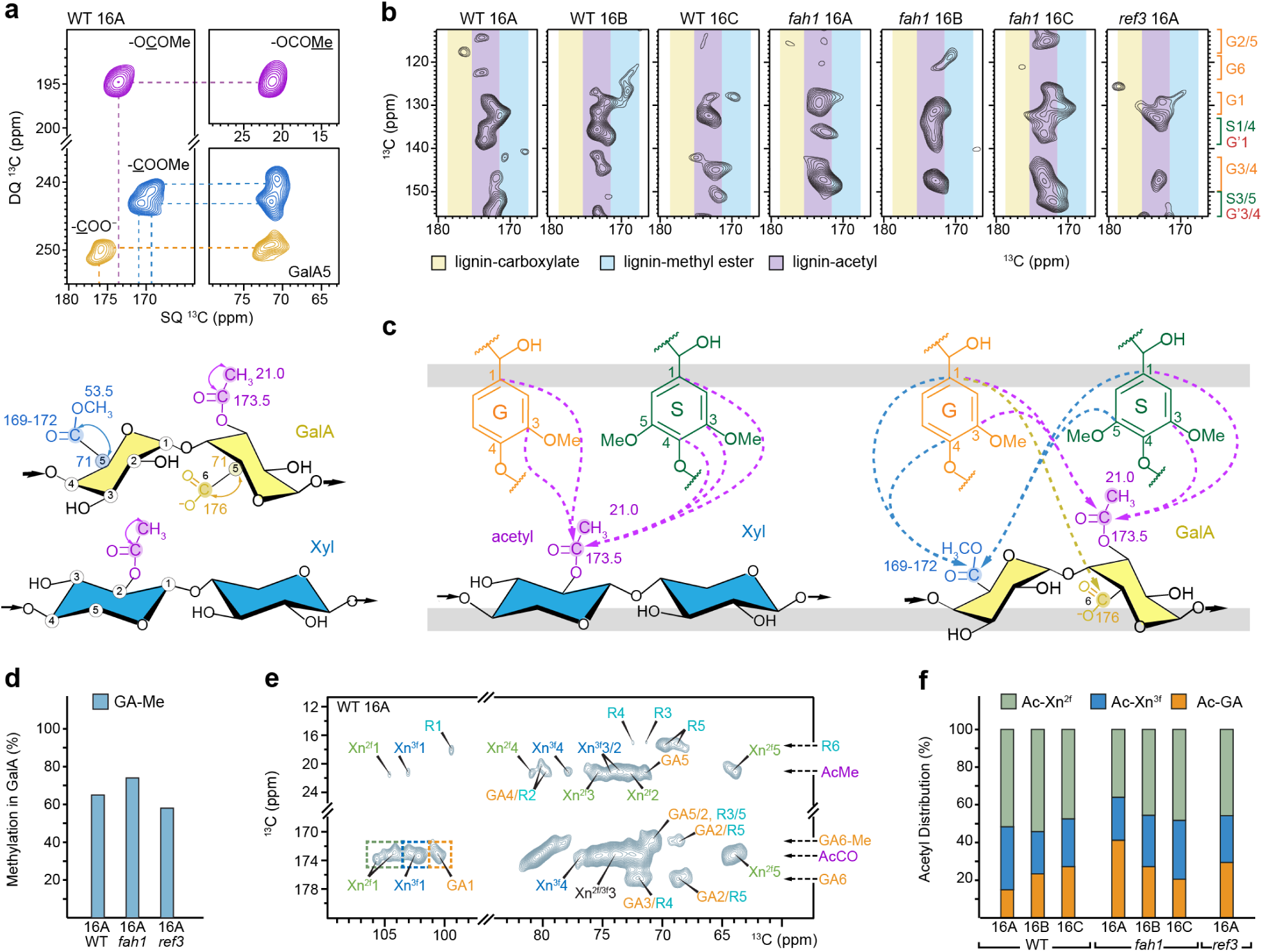
Pectin-lignin interactions identified in inflorescence stems. (**a**) Resolved signals of carbonyl sites from acetyl groups (-OCOMe, purple) present in both xylan and pectin, as well as methyl esters (-COOMe, blue) and carboxylates (-COO^-^, orange) in pectin, observed in CP-based refocused J-INADEQUATE spectra. The representative chemical structure and corresponding chemical shifts of these key carbon sites are also provided. (**b**) Specific cross-peak regions extracted from 2D dipolar-gated ^13^C-^13^C correlation spectra with a long mixing time of 1 s. The purple band highlights interactions between lignin aromatic carbons and acetyl groups. Yellow and blue bands indicate lignin interactions with pectin carboxylates and methyl esters, respectively. (**c**) Structural representation illustrating intermolecular interactions between lignin aromatic and carbonyl sites in xylan (blue) and the GalA units in HG (yellow). Note that GalA residues may also originate from RG-I. Black dashed lines represent intermolecular interactions shown in panel (b), while solid lines indicate intramolecular interactions observed in the spectra in panel (a). (**d**) Molar fraction of methylated GalA estimated by comparing the peak volumes of the resolved signals corresponding to methylated and unmethylated GalA units in spectra similar to the one shown in panel (a). (**e**) Cross peaks between methyl/carbonyl groups and other carbohydrate carbons observed in 2D dipolar-gated spectrum with a 0.1 s mixing time. NMR abbreviations are used for rhamnose (R), 3-fold xylan (Xn^3f^), 2-fold xylan (Xn^2f^), and galacturonic acid (GA). Dashed arrows highlight chemical shifts from the C6 of rhamnose (R6; 18.5 ppm), acetyl methyl (AcMe; 21 ppm), methylated GalA C6 (GA6^Me^; 171 ppm), acetyl carbonyl (AcCO; 173.5 ppm), and unmethylated GalA C6 (GA6; 176 ppm). Dashed boxes in green, blue, and orange highlight the integration regions used for estimating the distribution of acetyl groups in different carbohydrates. (**f**) Molar distribution of acetyl groups in Xn^2f^, Xn^3f^, and GalA, respectively.

These interactions are summarized in the structural model shown in **Fig. 6c**. It is evident that G-residues primarily interact with pectin via its methyl ester groups, with the carboxylate groups serving as secondary interaction sites. In contrast, S units did not exhibit any detectable interactions with pectin but were correlated with acetyl groups, which are likely mainly derived from xylan in these samples. Additionally, interactions were observed between lignin methoxy groups and the acetyl groups of both xylan and pectin, as well as the methyl esters of pectin, but not with the carboxylate groups (**Supplementary Fig. 16**). This supports the idea that unmethylated pectin seldom interacts with lignin, even during the early stages of lignification.

Echoing the NMR observations, Mäule staining of wild-type inflorescences also revealed clear localization of G-lignin in the middle lamella, situated between adjacent S-lignified fiber cells, and cell corner during the later stages of inflorescence maturation. The deposition of G-lignin was most prominent in the fully lignified basal region of 40 cm-tall wild-type stems (**Supplementary Fig. 4**) but was also detectable at an earlier developmental stage (34 cm). These fluorescence microscopy observations corroborate our NMR results, which indicate interactions between G-lignin and pectin, a key polymer of the middle lamella, aligning with previous reports of G-lignin accumulation in this region between adjacent cells^46,47^.

The pectic polymers in these *Arabidopsis* samples were found to be heavily methylated. The extent of methylation was estimated to be in the range of 65-77% for the basal segment of most WT samples and the *fah1* mutant, respectively (**Fig. 6d**; **Supplementary Fig. 9c**). Meanwhile, the *ref3* mutant showed a slightly lower degree of methylation at 58% (**Fig. 6d**), although the overall pectin content remained similar (**Supplementary Fig. 16**).

The next question is whether xylan or pectin predominantly contributed to the observed acetyl-lignin cross peaks. In dicot xylan, approximately 50% of xylosyl residues are *O*-acetylated at the C2 and/or C3 positions, which hinders enzymatic degradation and, when acetates are released, inhibits microbial fermentation during bioenergy production^25,48,49^. Similarly, *O*-acetylation occurs in pectin, where GalA residues in the backbones of homogalacturonan (HG) and rhamnogalacturonan I (RG-I) can be *O*-acetylated at the C2 and/or C3 positions (40-85% acetylation), and rhamnose (Rha) residues may also be acetylated at C3 sites^50–52^. We observed that the carbonyl carbon of acetyl groups (AcCO; 173.5 ppm) showed resolved correlations with carbon 1 of 2-fold xylan (Xn^2f^1), 3-fold xylan (Xn^3f^1), and GalA (GA1), enabling spectral deconvolution to estimate the molar distribution of acetyl groups between xylan and pectin (**Fig. 6e**). Consistently observed in most samples, 75-83% of acetyl groups were associated with xylan, while 15-27% were in pectin, with one exception in the *fah1-*16A sample, where 41% of acetyl groups were in pectin but still lower than in xylan (**Fig. 6f**). These findings further support the dominant role of xylan over pectin in mediating lignin interactions.

## DISCUSSION

Lignification is initiated at the cell corners and the pectin-rich middle lamella, suggesting a critical role for pectin in the early stages of this process^47,53^. It has been observed that pectin can coexist with lignin in the same polymer cluster and promote lignin polymerization during the biomimetic polymerization of coniferyl alcohol in pectin solutions^54^; however, direct evidence of pectin-lignin interactions *in muro* has not been observed previously. Our solid-state NMR evidence revealed that G-lignin and a minor proportion of H-units specifically interact with pectin (**Fig. 4c**, **Fig. 6b, c**), with these interactions being stronger in methylated pectin compared to unmethylated pectin. The degree of pectin methylation influences the formation of egg-box structures through Ca²⁺ chelation, alters the conformation of homogalacturonan (HG), and impacts cellulose organization, wall integrity, and mechanics^55–59^. Our data also reveals an unexpected role of pectin methyl esters in stabilizing interactions with deposited lignin during the early stages of lignification, where younger plants and segments exhibit relatively higher G-lignin and pectin content (**Fig. 1d, e**; **Supplementary Fig. 17**). Meanwhile, acetyl groups exhibit strong cross peaks with lignin, particularly with S-units (**Fig. 6b, c**); however, in the samples analyzed, acetyl groups are predominantly associated with xylan rather than pectin (**Fig. 6f**). These experimental findings highlight the distinct structural roles of S- and G-lignin in stabilizing different carbohydrate packing interfaces, likely linked to the temporal sequence of G- and S-rich lignin deposition, with the latter occurring at a later developmental stage as the plant deposits more secondary cell walls (**Fig. 1d**; **Supplementary Fig. 4**).

Previous solid-state NMR studies have shown that the strength of physical interactions between lignin and carbohydrates correlates with the molar fraction of S units in cell walls, particularly linked to the abundance of methyl substitutions in lignin across various plant species^28,29,60^. Such observations have emphasized the role of S-lignin in interacting with carbohydrates, particularly xylan, and the importance of electrostatic contacts in stabilizing the packing interface formed by the polar functionalities of xylan and lignin. The results reported in this study further confirmed this, as the low-S *fah1* mutant displayed the weakest lignin-carbohydrate cross-peak intensities (**Fig. 5a**) and the lowest mechanical strength and modulus (**Fig. 2f-h**). In contrast, the low lignin content of the *ref3* mutant did not affect mechanical properties, but the sparseness of G-lignin in this mutant promoted molecular mixing of remaining S-lignin with carbohydrates (**Fig. 5a**). These findings also provide two novel insights into the carbohydrate-lignin packing interface. First, molecular-level mixing, rather than lignin content alone, regulates the mechanical properties of secondary plant cell walls. This is consistent with the results of previous studies of woody tissues, which have indicated that lignin content primarily affects axial compression mechanics, whereas tensile strength correlates more closely with the cellulose microfibril angle^61,62^. Second, S-lignin alone, which is rich in methoxy groups, can effectively stabilize lignin-carbohydrate interactions when G-lignin is absent. This principle, identified through comparisons between WT and mutant plants, is further supported by the observation that older stem segments in WT plants exhibit higher levels of S-lignin and greater lignin-carbohydrate integration (**Fig. 5f**).

Despite these conceptual advances, many questions remain regarding the lignin-carbohydrate packing interface in lignocellulose. A key challenge is integrating the significance of lignin-carbohydrate electrostatic contacts observed through solid-state NMR spectroscopy with the covalent linkages previously identified in the lignin-carbohydrate complex (LCC)^63–68^. While these covalent linkages are essential for chemically stabilizing the polymer complex, they are relatively sparse, and the cell wall needs to rely on non-covalent interactions for overall structural integrity. In commelinid grasses, ferulate units link to arabinose residues on xylan sidechains, potentially facilitating cross-linking with bulk lignin domains through radical coupling, as observed in in-vitro lignification and LCC analysis^69–72^. A recent solid-state NMR study in *Brachypodium* highlighted the role of such linkages, showing that ferulated and acetylated xylan chains with a two-fold conformation extensively interact with cellulose in *Brachypodium* stems, forcing lignin to bind with cellulose microfibrils associated with these xylan chains^28^. In *Arabidopsis* secondary cell walls, glucuronic acid (GlcA) sidechains on glucuronoxylan likely mediate xylan-lignin cross-linking, and the removal of GlcA units in an *Arabidopsis gux* mutant decreases recalcitrance and increases xylose release by 7-fold during saccharification^73,74^. However, the available evidence for such linkages in *Arabidopsis* is indirect^74^. The structural significance of this proposed cross-linking mechanism to the cell wall architecture of *Arabidopsi*s and hardwood lignocellulose requires further investigation.

Efforts to detect interactions between lignin and the GlcA sidechains of xylan in *Arabidopsis* were unsuccessful. In *Arabidopsis*, GlcA branches are sparsely distributed, with approximately one in eight xylose residues carrying a GlcA substitution via α-1-2 linkage, and the majority of these GlcA units are heavily methylated at the O-4 position (4-O-MeGlcA) with a GlcA to 4-O-MeGlcA ratio of one to three^25,75–77^. GlcA in xylan is extensively 4-O-methyl-etherified, with a distinct chemical shift of 58.5-60.5 ppm, differing from the 53.5 ppm 6-O-Me signal of GalA in pectin^78–80^. Additionally, the C1 chemical shifts for GalA are resolved and distinguishable from the mixed signals of GlcA and xylose sidechains in xyloglucan (**Supplementary Fig. 18**). Despite these resolved signals, no lignin-GlcA contacts were observed in the 2D ^13^C-^13^C correlation spectra (**Supplementary Fig. 19**). This could stem from the low abundance of GlcA in *Arabidopsis* cell walls, where GlcA constitutes less than 4% of non-crystalline polysaccharides^81^, while GalA is five times more abundant, or the infrequency of lignin-carbohydrate covalent linkages in the cell wall. While these putative linkages may act as anchors between lignin and xylan, the stability of the cell wall architecture is likely maintained predominantly by physical interactions.

## METHODS

### Growth and harvest of *Arabidopsis* stem material

Wild-type (Col-0) *Arabidopsis* plants were grown hydroponically according to previously published methods, with some modifications. Black polycarbonate food pans (Sparrow Food Solutions; Sixth Size Polycarbonate Food Pan - 2.5" Deep, PLPA8162/BK) were filled to the brim (∼800 mL) with “Standard Solution” hydroponic media^82^. Five 8-mm holes were drilled through the pan lids (Sparrow Food Solutions; Black Polycarbonate Food Pan Lids, THPLPA7160CBK) at a center-to-center distance of approximately 5 cm. 0.65 mL Eppendorf tubes were filled to the top, forming a bubble over the opening of the tube, with 0.35% agar (Sigma; A7921) solution (dissolved in ddH_2_O by boiling). Once solidified, the bottom, tapered part of each tube was cut off with a razor blade. The tubes were placed in the food pan lid holes with wire mesh to prevent the agar from slipping out of the tube^82^. The lid with suspended tubes was placed on top of the pan containing hydroponic media, ensuring that each tube was fully submerged in the nutrient solution to maintain hydration.

*Arabidopsis* seeds were sterilized and germinated directly on the agar in Eppendorf tubes, suspended in hydroponic nutrient media. Plants were grown on a 12-hour day (22°C)/12-hour night (16°C) cycle with 120 µmol/m^2^/s white light. Inflorescence stems began to emerge from rosettes six to eight weeks after germination. Nutrient solution was replaced as needed to ensure the base of each tube remained submerged. Inflorescence stems were measured and cut away from the rosette at pre-determined heights, cut into 2 cm segments from the base (basal) to the top (apical) of stem, and immediately frozen at −80°C.

^13^C-labeling of inflorescence stems was achieved by hydroponic growth of stems in a sealed chamber (60 cm x 42 cm x 52 cm), fashioned from Plexiglass with butyl rubber weather stripping to ensure it was airtight. ^13^C-labled carbon dioxide (Sigma-Aldrich 364592) was dosed into the chamber by a digital CO_2_ controller (Titan Controls HGC702853) and was maintained at 600ppm, beginning when inflorescences were just emerging (0 cm) to harvest at the pre-specified heights. Ethylene gas absorber packets (Fresh & Fresh) and indicating desiccant chips (DRIERITE 21001) were used to control ethylene and the humidity within the chamber, and were swapped out as needed to maintain the optimal growth of plants in the sealed environment.

### Mäule Staining and microscopic observation of lignin distribution

*Arabidopsis* stems were harvested at predetermined heights and the segments of interest were excised and submerged in cryomatrix (Tissue-Tek® OCT Compound; Ted Pella 27050) for 48 hours. Segments were then positioned in molds and frozen at −80°C in cryomatrix until the time of sectioning. 50 µm cross sections were made with a Leica CM1950 cryostat and placed on silane-coated glass slides. The sections were washed in ddH_2_O, shaking at 4°C overnight. Sections were then cleared with ClearSee solution^83^ at 22°C without shaking for 24 hours with one exchange of ClearSee solution. Cleared sections were washed with ddH_2_O to remove ClearSee and then Mäule stained^84,85^. Stem cross sections were stained for 5 minutes in 1% (w/v) potassium permanganate (KMnO4) solution. Stained sections were rinsed with ddH_2_O to remove residual KMnO_4_ and then incubated in 1N HCl for 5 minutes. HCl solution was removed and sections were rinsed with 1M Tris-HCl buffer (pH 8) and mounted in the same buffer with a glass coverslip for microscopic observation. A Zeiss LSM780 confocal microscope was used to observe staining in stem cross sections. G lignin was observed with 488 nm excitation and 510-580 nm emission. S lignin was observed with 488 nm excitation and 600-660 nm emission. All micrographs were captured with the same imaging settings.

### Mechanical analysis of inflorescence stem segments

Frozen 3-cm basal stem segments from 16 cm tall inflorescences were thawed and flattened for 5 seconds with 60 psi of pressure between paddle jaw faces (Intron 2702-375) on a tensile stretching device (Instron 68SC-05). After flattening, the epidermis was removed from the stem segments by peeling with forceps. Stem segments were clamped with rubber-coated jaw faces (Instron 2702-360) with added sandpaper for extra grip, at a 6 mm clamping distance. Stems were stretched with monotonic loading to failure. Eight basal stem segments (biological replicates) were analyzed per genotype. The stress-strain curves were plotted with Python. The strain was calculated as the displacement divided by the original sample length. The stress was determined by dividing the force by the estimated cross-sectional area, calculated as the cell wall mass divided by the product of cell wall density (previously reported as 1.5 g/cm³)^86,87^ and stem length. To measure the cell wall mass, the 6 mm stem segment between the clamps was excised and chemically and enzymatically treated to remove non-cell wall components. Stem segments were incubated in 1.5% sodium dodecyl sulfate (SDS) at 22°C, shaking for 16 hours to remove proteins. The tissue was then washed 10 times with double-distilled water (ddH_2_O). Stems were then incubated at 37°C for 24 hours, shaking, in 50 mM MES buffer (pH 6.8) that contained porcine pancreas α-amylase (5000 units/30 mL) to remove starch and 0.02% NaN_3_ to inhibit microbial growth. Finally, the tissues were washed again in 1.5% SDS at 22°C for 1 hour and rinsed 10 times in ddH_2_O with 0.02% NaN_3_, and oven-dried at 65°C for 72 hours. Finally, the treated, dried stem segments were weighed.

### Preparation of *Arabidopsis* samples for solid-state NMR analysis

Three genotypes of *Arabidopsis thaliana* (Columbia-0) were analyzed: the wild-type (WT), the *fah1-2* mutant (ferulic acid 5-hydroxylase 1)^37,38,41^, and the *ref3* mutant (*ref3-3*, reduced epidermal fluorescence 3-3)^34–36^. Six distinct growth stages of WT stems were examined, characterized by total plant heights of 8, 12, 16, 19, 24, and 32 cm. The growth stages at 8, 12, and 16 cm consistently achieved their specified heights. However, the latter stages—19, 24, and 32 cm— exhibited variability in their final heights, and the average height within the respective ranges was reported. Specifically, the 19, 24, and 32 cm stages corresponded to height ranges of 18–20 cm, 23–25 cm, and 30–34 cm, respectively, as illustrated in **Supplementary Figure 20d–f**. For the *fah1* and *ref3* mutants, only the 8 and 16 cm growth stages were investigated.

The inflorescence stems used for NMR analysis were segmented into three regions: basal, middle, and apical. The basal region was further subdivided into three distinct segments: A (0–2 cm), B (2–4 cm), and C (4–6 cm). The cutting scheme for basal segments A/B/C was consistent across all samples. However, the cut regions for the middle and apical sections varied depending on the growth height. A visual representation of the cutting method is provided in **Supplementary Figure 20**, and a comprehensive summary of all cut regions and corresponding samples used for NMR analysis is presented in **Supplementary Table 2**. For each NMR experiment, inflorescence stems from multiple plants of the same growth stage were collected. The same segment was consistently harvested across stems from a given growth stage. The collected material was combined and shredded into small fragments before being loaded into the NMR rotor. The stems used for analysis were never dried to preserve their native state.

### Solid-state NMR experiments

Solid-state NMR experiments were conducted using three Bruker Avance Neo spectrometers equipped with 3.2-mm MAS triple resonance probes (^1^H/^13^C/^15^N). The spectrometers operated at three magnetic field strengths: 400 MHz (9.4 Tesla) and 600 MHz (14.1 Tesla) at Michigan State University, and 700 MHz (16.4 Tesla, Bruker Ascend) at Louisiana State University. Experiments were performed with MAS spinning rates between 14-20 kHz at sample temperatures ranging from 275 K to 298 K. ^13^C chemical shifts were externally referenced to tetramethylsilane (TMS) by calibrating the adamantane CH₂ resonance peak to 38.48 ppm, with the resulting spectral reference (sr) applied to the plant sample spectra. Radiofrequency field strengths were typically 83 kHz for ^1^H decoupling, and 83.3 kHz and 50–62.5 kHz for the 90° hard pulses of ^1^H and ^13^C, respectively. The key parameters and conditions of NMR experiments are listed in **Supplementary Table 3**.

1D ^13^C NMR spectra were acquired using different polarization methods to investigate the structure and dynamics of molecules. ^1^H-^13^C cross-polarization (CP) experiments were measured on all samples to selectively detect rigid molecular components. CP transfer efficiency is affected by factors such as the protonation state and dynamics, therefore compositional analysis of this experiment only serves as an estimate rather than quantification. Meanwhile, quantitative compositional analysis was achieved using direct polarization (DP) with a long (35 s) recycle delay that allowed complete relaxation of all ^13^C magnetization to equilibrium between scans, enabling the quantitative detection of all carbons in the sample. This method was employed for most samples, with the exception of WT08A, WT12A, *fah1-*08A, and *ref3-*08A. In parallel, to specifically target mobile molecules, the same DP scheme was coupled with a short (2 s) recycle delay that allowed only the relatively dynamic molecules, characterized by fast ^13^C-T_1_ relaxation, to relax to equilibrium between scans, enabling selective detection of these mobile components. This approach was applied to samples WT16A/B/C, *fah1-*16A/B/C, and *ref3-*16A.

All 2D NMR spectra were acquired using CP-based methods with phase-sensitive quadrature detection (States acquisition)^88^. Chemical shift assignment was performed using DQ-SQ correlation spectroscopy, employing the refocused Incredible Natural Abundance DoublE QUAntum Transfer Experiment (INADEQUATE) pulse sequences^89,90^. The DQ-SQ correlation approach enhances spectral resolution by suppressing diagonal peaks and spreading chemical shift information into the indirect DQ dimension. In this dimension, the chemical shift corresponds to the sum of the isotropic chemical shifts of a coupled spin pair, providing improved peak separation for the measured spectrum. The coherence transfer in the INADEQUATE sequence is mediated through J-coupling, ensuring that the observed correlations are exclusively between directly bonded spin pairs. This feature enables unambiguous determination of carbon connectivity within the sample. The ^13^C chemical shifts are documented in **Supplementary Table 4**. CP-based J-INADEQUATE spectra were acquired for samples WT16A, WT19A, WT32A, *fah1-*16A, and *ref3-*16A.

To enhance the detection of lignin signals, which are typically masked by the strong carbohydrate signals, an aromatic editing technique was employed^91^. This approach incorporates a dipolar-gating period that suppresses signals from protonated carbon sites and attenuates those from proton-rich carbohydrates, while minimally affecting deprotonated carbon sites such as specific aromatic carbons in lignin (**Supplementary Fig. 21**)^91^. In conjunction with the dipolar-gating method, two homonuclear recoupling techniques were used to establish through-space correlations: Proton-Driven Spin Diffusion (PDSD) and Dipolar-Assisted Rotational Resonance (DARR)^92^. DARR experiments utilized a mixing period of 100 ms with low-power proton irradiation, optimizing the detection of intramolecular interactions within a distance of up to 6 Å. In contrast, PDSD experiments, which do not require proton irradiation, employed a longer mixing period of 1 s to capture relayed spin transfers, enabling the detection of long-range intermolecular interactions up to 10 Å^93,94^. Gated-DARR and PDSD spectra were collected for samples WT08A, WT12A, WT16A/B/C, *fah1-*16A/B/C, and *ref3-*16A, providing detailed structural insights into lignin and its spatial interactions with surrounding components.

### Estimation of molecular composition

Most spectra were processed using Bruker TopSpin software (version 4.3.0) with Lorentzian-to-Gaussian apodization functions unless specified otherwise. Detailed processing parameters for each spectrum are provided in **Supplementary Table 3**. For the analysis of 1D spectra, four integration regions (A1–A4) were defined and are depicted in **Supplementary Fig. 5**. Region A1 (190–0 ppm) encompasses all ^13^C signals in the system. This range represents all rigid carbons in plant cells for 1D CP spectra, all carbons in plant cells for 1D DP spectra, and all carbon signals propagated from the source carbon site in the indirect dimension (F1) for 1D slices extracted from 2D spectra. Region A2 (106–59 ppm) includes carbohydrate ring carbons. While minor overlap occurs between the lignin S2/6 peaks and carbohydrate signals such as Xn and cellulose C1 (106– 102 ppm), their contributions to A2 are negligible due to the carbohydrate signals being at least an order of magnitude stronger than those of lignin. Region A3 (156–109 ppm) represents the primary aromatic carbon region. It captures lignin aromatic carbons, excluding S2/6, as well as potential contributions from proteins and lipids. Region A4 (156–140 ppm) is uniquely assigned to lignin, representing a set of peaks that do not overlap with signals from proteins or lipids. The integration regions and their corresponding carbon species assignments are summarized in **Supplementary Table 5**.

The rigid lignin content in the whole cell is estimated based on the percentage of the ^13^C signal, calculated as the ratio of integrations A3/A1 from the 1D CP spectra. While the DP method provides a more quantitative analysis of the composition, the A3 region in DP spectra contains significant contributions from proteins and lipids, in addition to lignin signals. Notably, strong aromatic residue sidechain and fatty acid chain peaks appear at 136, 130, and 128.5 ppm in this region. In contrast, the A4 region in both CP and DP spectra is free from protein and lipid contributions, containing exclusively lignin signals. The line shape in this region remains consistent between the CP and DP spectra (**Supplementary Fig. 5c**). To reconcile these differences, we assume that the A4/A3 ratio is the same for both CP and DP spectra. This ratio is then used as a scaling factor to estimate the overall lignin content in the 1D DP spectra. Consequently, the quantitative lignin content is calculated by dividing the A4/A1 ratio in the DP spectrum by the A4/A3 ratio obtained from the CP spectrum.

### Analysis of lignin-carbohydrate packing interactions

For the analysis of long-range interactions in the 2D gated-PDSD spectra, eight integration regions (B1–B8) were defined. Region B1 contains all ^13^C signals propagated from deprotonated lignin carbons, specifically from S-lignin carbons at positions 1, 3–5, and G-lignin carbons at positions 1, 3, and 4. Region B2 encompasses ^13^C signals propagated from methoxy group (OMe) carbons, primarily from the methyl ether groups of lignin (∼56.5 ppm), as well as a minor contribution from methyl esters (-COOMe) of methylated galacturonic acid (GalA) in pectin (∼53.5 ppm). Region B3 includes ^13^C signals propagated from carbohydrates, originating from deprotonated lignin carbons. Similar to the A2 region in the 1D spectra, this region contains a small contribution from the S2/6 peaks, but these contributions are negligible compared to the stronger carbohydrate signals. Region B4 consists of carbohydrate ^13^C signals propagated from OMe carbons. Region B5 contains carbonyl (CO) ^13^C signals propagated from deprotonated lignin carbons, including signals from the galacturonic acid (GalA) carboxylate carbon (-COO⁻, ∼176 ppm), the acetyl carbonyl (AcCO, ∼174 ppm) from either Xn or GalA, and the carbonyl signals from methylated GalA (-COOMe, 172–168 ppm). Region B6 includes acetyl methyl (AcMe) group ^13^C signals, either from Xn or GalA, propagated from deprotonated lignin carbons. Region B7 captures all carbonyl (CO) signals propagated from the OMe groups. Finally, region B8 contains all AcMe signals propagated from the OMe groups.

Long-range interactions in the 2D gated-PDSD spectra are mediated by ^13^C polarization transfers within a spin diffusion range of approximately 1 nm. The strength of these interactions is quantified as the percentage of specific cross-peak signals (sink) relative to all ^13^C signals from the same source, and is calculated based on peak integration ratios^29,93^. For instance, the strength of the OMe-carbohydrate (source-sink) interaction is calculated as the ratio of the integration of region B4 (sink) to region B2 (source). The interaction strength, derived from the 1D cross-section of the 2D gated-PDSD spectrum, follows a similar calculation method using the integration regions defined in the 1D analysis described earlier. For example, the OMe–carbohydrate interaction strength can be calculated from the 1D slice as the ratio of integration area A2 (sink) to A1 (source). The integration regions, types of interactions, and detailed calculation methods are summarized in **Supplementary Table 5**.

### Analysis of carbohydrate acetylation and methylation and lignin S/G ratio

To analyze the content of acetylation in different polysaccharides, three integration areas of the 2D gate-DARR spectra were defined for the cross-peak regions corresponding to Xn^2f^, Xn^3f^, and GalA (**Fig. 6e**). The C1 sites of these three regions exhibit distinguishable chemical shift ranges, and their cross-peaks are well-resolved in most spectra, allowing their C1 sites to be used for the analysis. The AcCO-Xn^2f^, AcCO-Xn^3f^, and AcCO-GalA cross-peaks are used to represent the level of acetylation. The integration regions for these cross-peaks are summarized in **Supplementary Table 5**.

The level of GalA methylation was analyzed using 2D refocused J-INADEQUATE spectra. The C5-C6 spin pair of GalA was specifically used for methylation analysis, as these carbons have distinct chemical shift regions that do not overlap with other carbon sites in the DQ-SQ type spectrum. While the C5 chemical shift (∼72 ppm) is similar for both unmethylated GalA and methylated GalA (GalA-Me), their C6 chemical shifts differ: 176 ppm for unmethylated GalA and 168–171 ppm for methylated GalA. Therefore, only the integrations of the C6 cross-peaks were used to estimate the level of GalA methylation.

The S/G ratio of lignin was also estimated using 2D refocused J-INADEQUATE spectra. The G units exhibit a prominent cross-peak region, with single-quantum (SQ) chemical shifts ranging from 142-154 ppm and DQ chemical shifts between 290-300 ppm. This cross-peak corresponds to the G3-G4 spin pair, and its integration is used to estimate the G unit content. For the S lignin units, two prominent cross-peaks are observed, corresponding to the spin pairs S3/5-S4, with their SQ chemical shifts at approximately 154 ppm and 134 ppm, respectively, and a DQ shift around 288 ppm. Since these peaks represent two spin pairs, S3-S4 and S5-S4, which contribute equally, the S unit content is estimated by integrating both peaks and dividing the total by two.

## DATA AVAILABILITY

All relevant data that support the findings of this study are provided in the article and supplementary Information. All the original ssNMR data files will be deposited in the Zenodo repository and the access code and DOI will be provided. Source data will be provided as a Source Data file.

## AUTHOR CONTRIBUTIONS

P.X., D.D., and W.Z. performed solid-state NMR experiments. S.A.P. prepared ^13^C-labeled materials and conducted imaging analysis. C.-J.L. advised on the choice of the mutants and provided the mutant lines. D.C. and T.W. designed the experiments. All authors contributed to the data analysis and writing of the manuscript.

## ACKNOWLEDGMENT

The solid-state NMR analyses were supported by the U.S. Department of Energy under grant no. DE-SC0023702 to T.W. Sample preparation, imaging analysis, and mechanical measurements were supported as part of the Center for Lignocellulose Structure and Formation, an Energy Frontier Research Center funded by the US Department of Energy, Office of Science, Basic Energy Sciences under award no. DE-SC0001090. C.-J.L. was supported by the U.S. Department of Energy, Office of Science, Office of Basic Energy Sciences under contract number DE-SC0012704-specifically through the Physical Biosciences program of the Chemical Sciences, Geosciences and Biosciences Division. The authors thank Dr. Fabien Deligey for the initial NMR analysis, Dr. Jingyi Yu for assistance with mechanical analysis, and Andrea R. Brown for preparing the cartoons of Arabidopsis inflorescence stems.

