## Supplementary Information for "Emergence of Lignin-Carbohydrate Interactions During Plant Stem Maturation Visualized by Solid-State NMR"

This file includes:

Supplementary Figures 1-21 and Tables 1-5

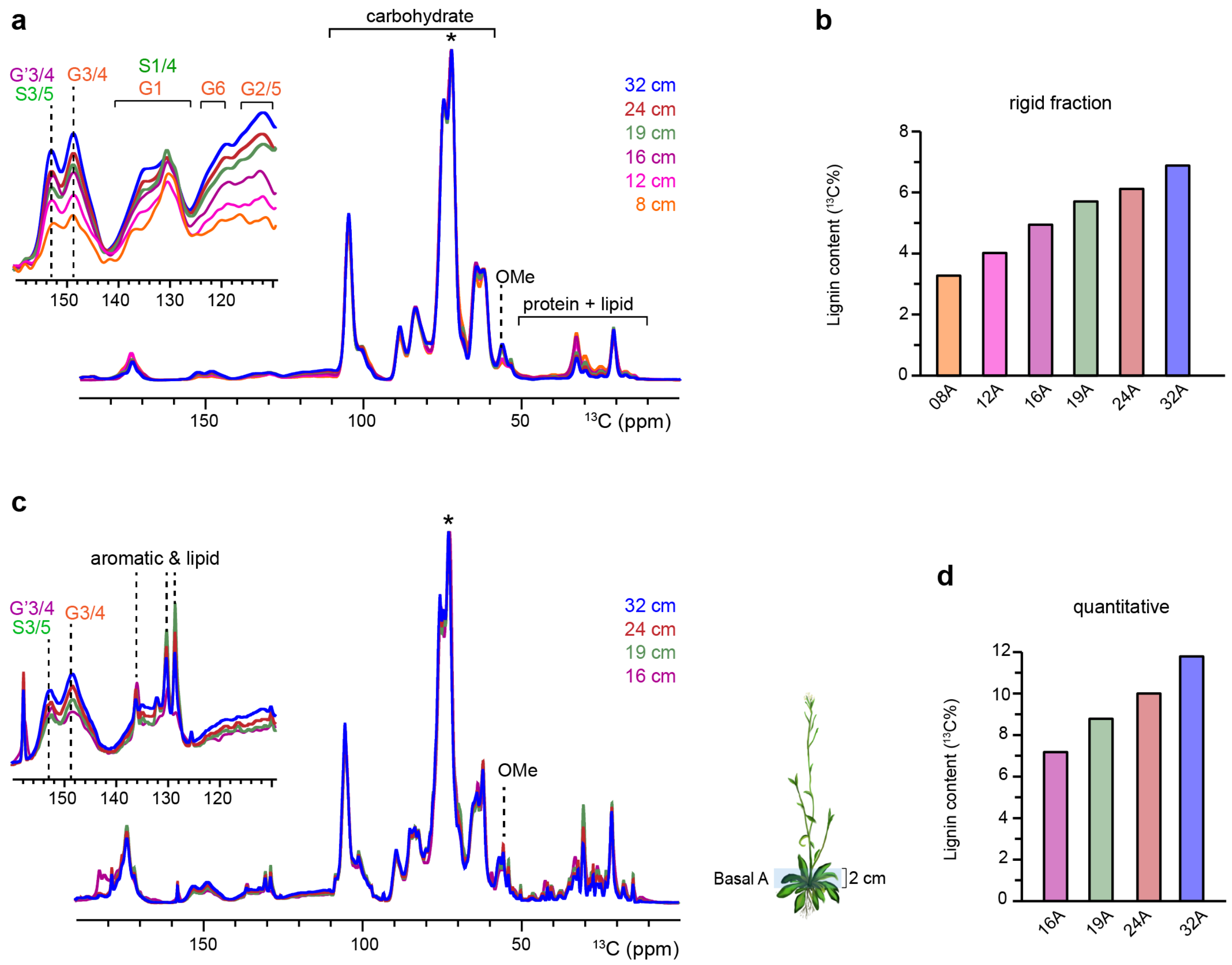


**Supplementary Figure 1. Lignification of WT *Arabidopsis* stems at various growth stages.** (**a**) 1D CP spectra showing rigid components of the WT *Arabidopsis* samples. All spectra are normalized with respect to the highest carbohydrate peak (asterisk). Zoomed-in view was provided for the lignin aromatic region showing the changes in the lignin content. The samples include the basal A of the stem grew to heights of 8, 12, 16, 19, 24 and 32 cm. (**b**) Overall lignin content in the rigid fraction of whole cell estimated from the 1D CP spectra of each sample. (**c**) 1D $^{13}$C DP spectra with long recycle delays of 35 s showing quantitative $^{13}$C composition of the WT *Arabidopsis* samples. The samples include the basal A of the stem grown to heights of 16, 19, 24 and 32 cm. The illustrations on the right show the inflorescence segment (basal A, 0-2 cm) used for all experiments. All spectra are normalized with respect to the highest carbohydrate peak (asterisk). Zoomed-in view region includes signals from the lignin aromatic carbons, fatty acyl chain carbons in lipids, as well as sidechain carbons in aromatic amino acid residues. The aromatic residues and lipids here are more dynamic and their signals manifest as strong sharp peaks at around 136, 132, 130 and 128 ppm. The unambiguous lignin peaks are at around 152.5 ppm (S3/5, G'3/4) and 148.5 ppm (G3/4); their peak shapes and positions are largely invariant compared to their counterparts in the CP spectra. (**d**) Quantitative analysis of overall lignin content in whole cell from $^{13}$C DP spectra of each sample.

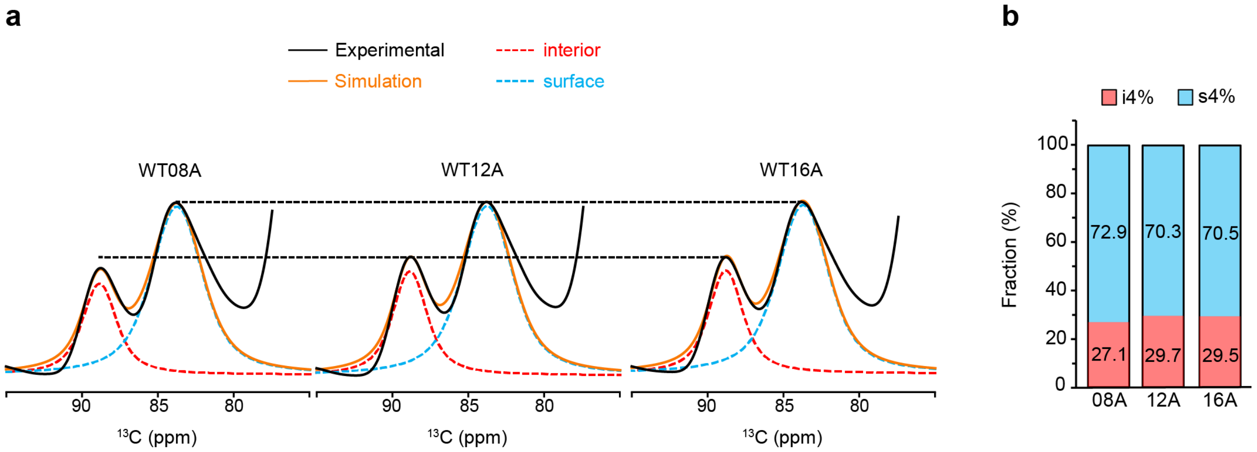


**Supplementary Figure 2. Estimation of surface and interior chains of cellulose in wild-type stems.** (**a**) Spectral deconvolution of the unambiguous cellulose region in 1D CP spectra for samples WT-08/12/16A. The resolved peaks for interior and surface cellulose are i4 (89 ppm) and s4 (84 ppm), respectively, and all spectra were normalized to the s4 peak. The experimentally measured spectra (black) are overlaid with the simulated spectra (orange) reconstructed from the deconvoluted peaks. The deconvolution was performed with ssNake software[1], using Lorentzian/Gaussian model. (**b**) Fractions of interior and surface cellulose chains in each sample. The percentages were estimated from the integrals of each deconvoluted peak of i4 and s4 and were indicated on each bar.

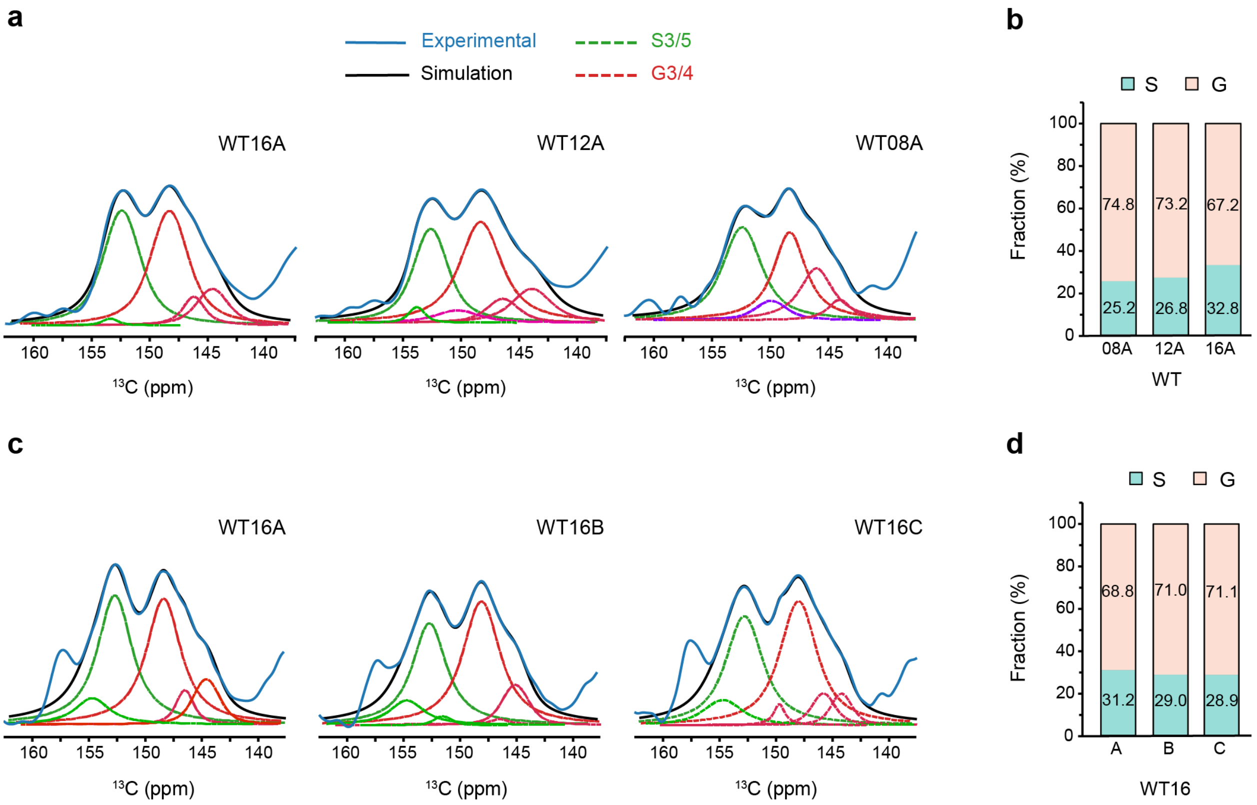


**Supplementary Figure 3. Estimation of S and G content in wild-type stems.** (**a**) Spectral deconvolution of the signature lignin region in 1D CP spectra for samples WT-08/12/16A. The experimentally measured spectra (blue) are overlaid with the simulated spectra (black) reconstructed from the deconvoluted peaks. The G units have many forms, some have extremely broad peaks (purple) extending across the S unit region, even after deconvolution. (**b**) Estimation of the fractions of the S and G unit in each sample. Estimations were based on the integrals of deconvoluted peaks of each unit, and the percentages were indicated on each bar. (**c**) Spectral deconvolution of 1D CP spectra for samples WT-16A/B/C. The peak at 157 ppm is the spinning sideband (SSB) artifact. (**d**) Estimation of the fractions of the S and G unit in each sample. All deconvolutions were performed with ssNake software, using Lorentzian/Gaussian model.

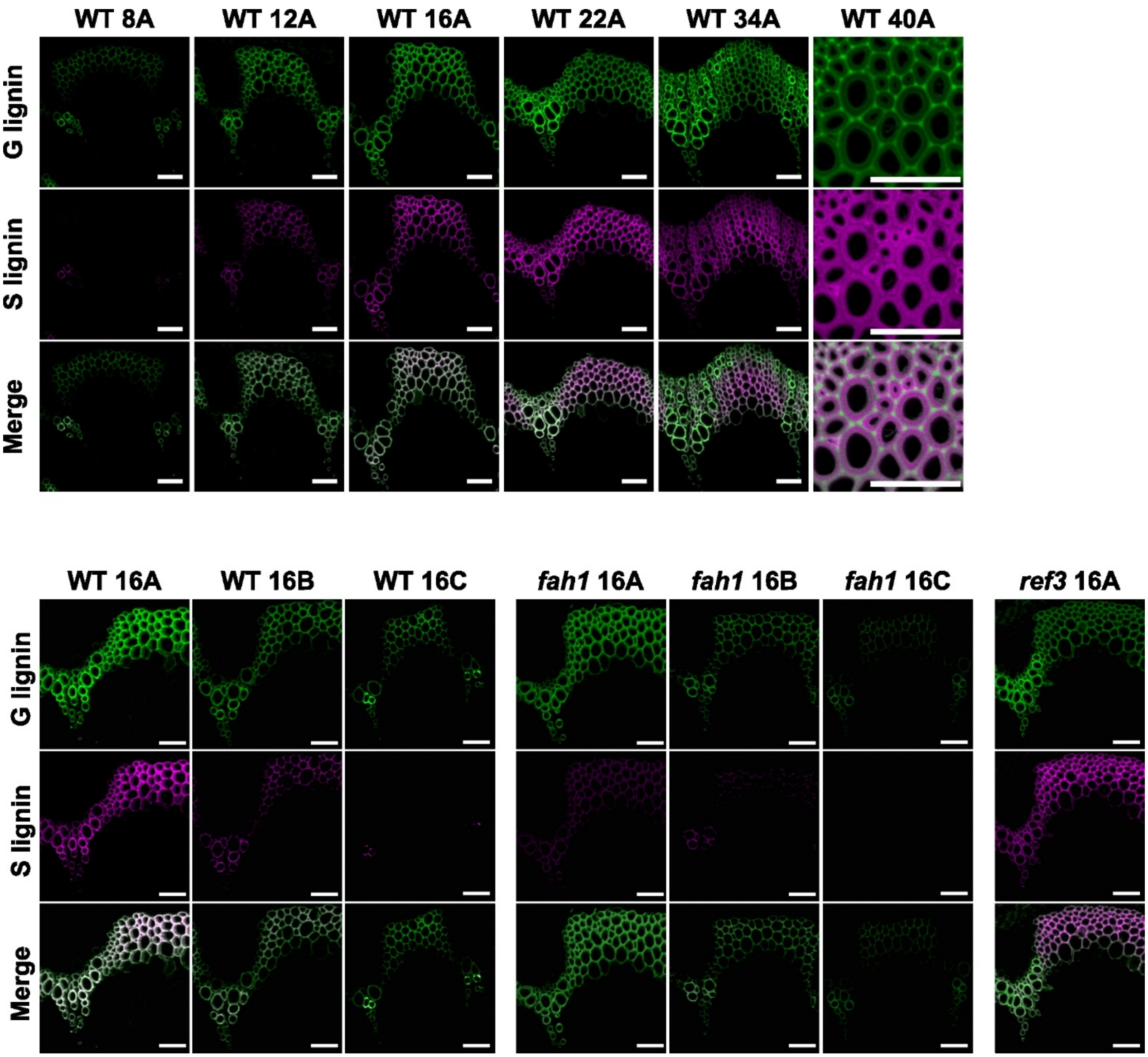


**Supplementary Figure 4. Single channel and overlay images of representative Mäule-stained *Arabidopsis* stem cross sections.** Fluorescence images depict the distribution of G-lignin (green) and S-lignin (purple) in the lignified tissues of basal segments of inflorescences across a developmental gradient in wild-type (top) and in two lignification mutants at SCW maturation (bottom). Scale bars = 50 µm.

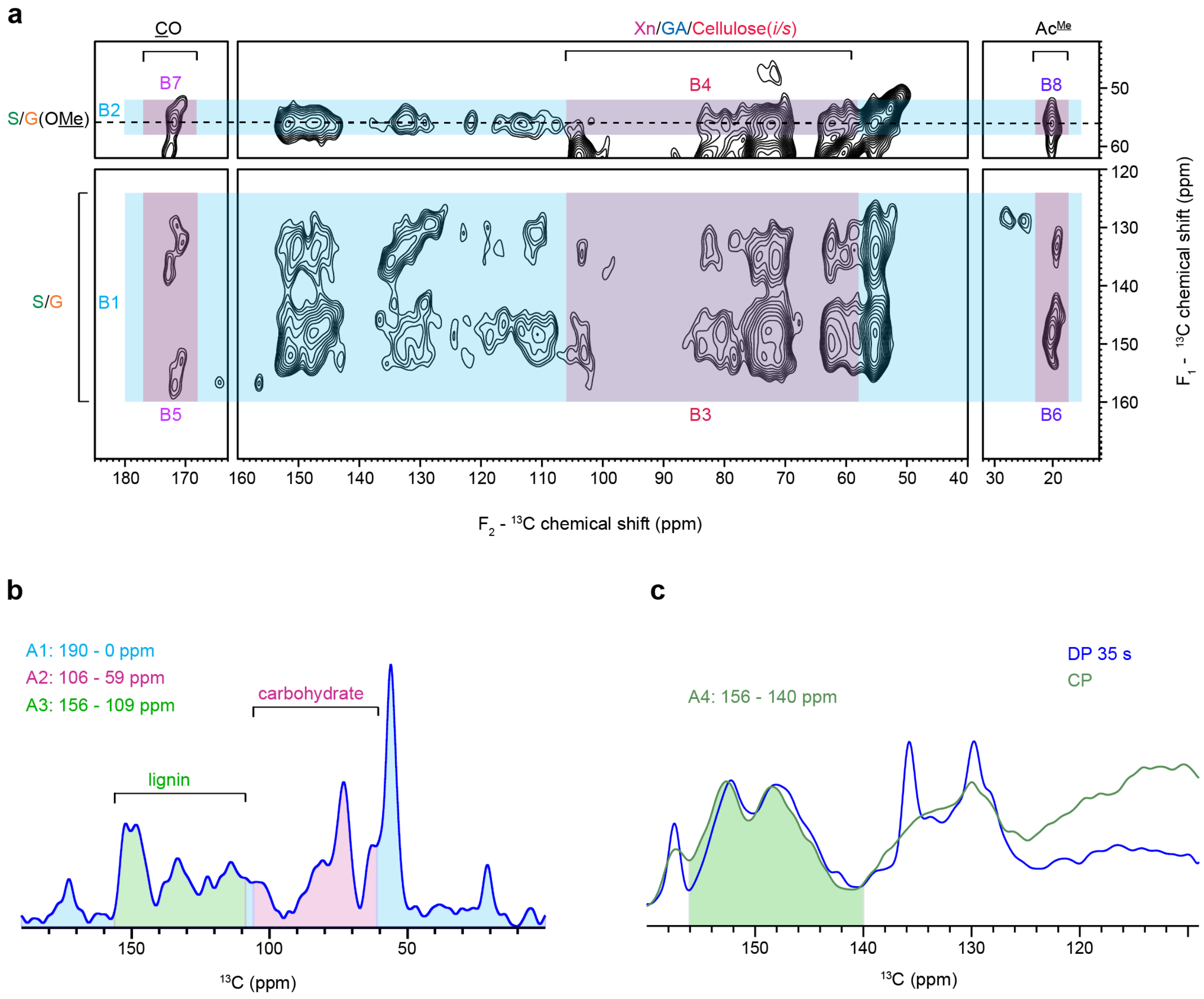


**Supplementary Figure 5. Integration areas used for analyzing lignin content and interactions.** (**a**) Integration areas (B1-B8) used for the analysis of 2D dipolar-gated PDSD spectra (1 s mixing time, long-range). Areas highlighted in blue indicate all $^{13}$C signals originated from lignin ring carbons and OMe carbons. The OMe carbons are mainly from the lignin methyl ether (56 ppm, indicated by horizontal dashed line) and may include minor pectin methyl ester contribution (53 ppm). Areas highlighted in dark magenta indicate cross-peaks between lignin aromatic carbons and carbohydrate carbons. (**b**) 1D cross section extracted from the 2D spectrum. Area A1 represents all $^{13}$C signals in the system. A2 includes carbohydrate signals. A3 includes lignin aromatic carbon sites aside from S2/6. The signals in A3 region are primarily from lignin in the CP experiment but also contain contributions from proteins and lipids in the DP experiment. (**c**) Zoomed-in view of the aromatic region with both the CP spectrum (green) and DP spectrum (blue). A4 indicates the unique lignin peak region used for the quantitative lignin content analysis. The line shape of this region is invariant between the CP and DP spectrum. The chemical shift ranges for each area are summarized in **Supplementary Table 5**.

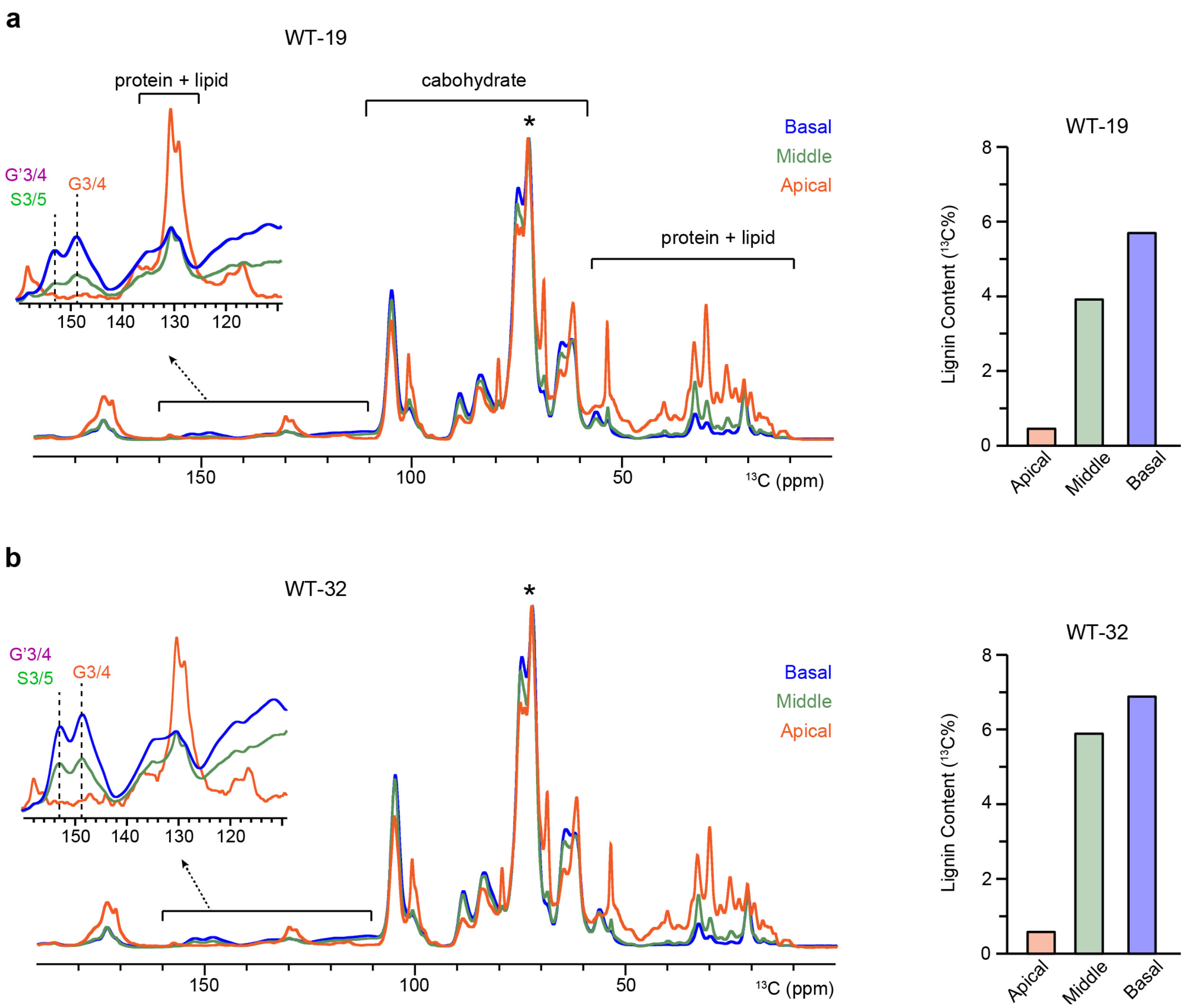


**Supplementary Figure 6. Lignification of WT *Arabidopsis* across the whole inflorescence.** 1D CP spectra collected from three different inflorescence regions were compared for the WT *Arabidopsis* grew to heights of 19 cm (**a**) and 32 cm (**b**). All spectra are normalized with respect to the highest carbohydrate peak (asterisk). The zoomed-in region shows the difference in the overall lignin content between different inflorescence regions. For both 19 cm and 32 cm samples, the unique lignin peaks at 152.5 ppm (S3/5, G'3/4) and 148.5 ppm (G3/4) are depleted in the apical segment (orange), whereas the aromatic residue and lipid peaks became prominent and dominant. The overall lignin content in the rigid fraction of whole cell estimated from the 1D CP spectra are shown on the right.

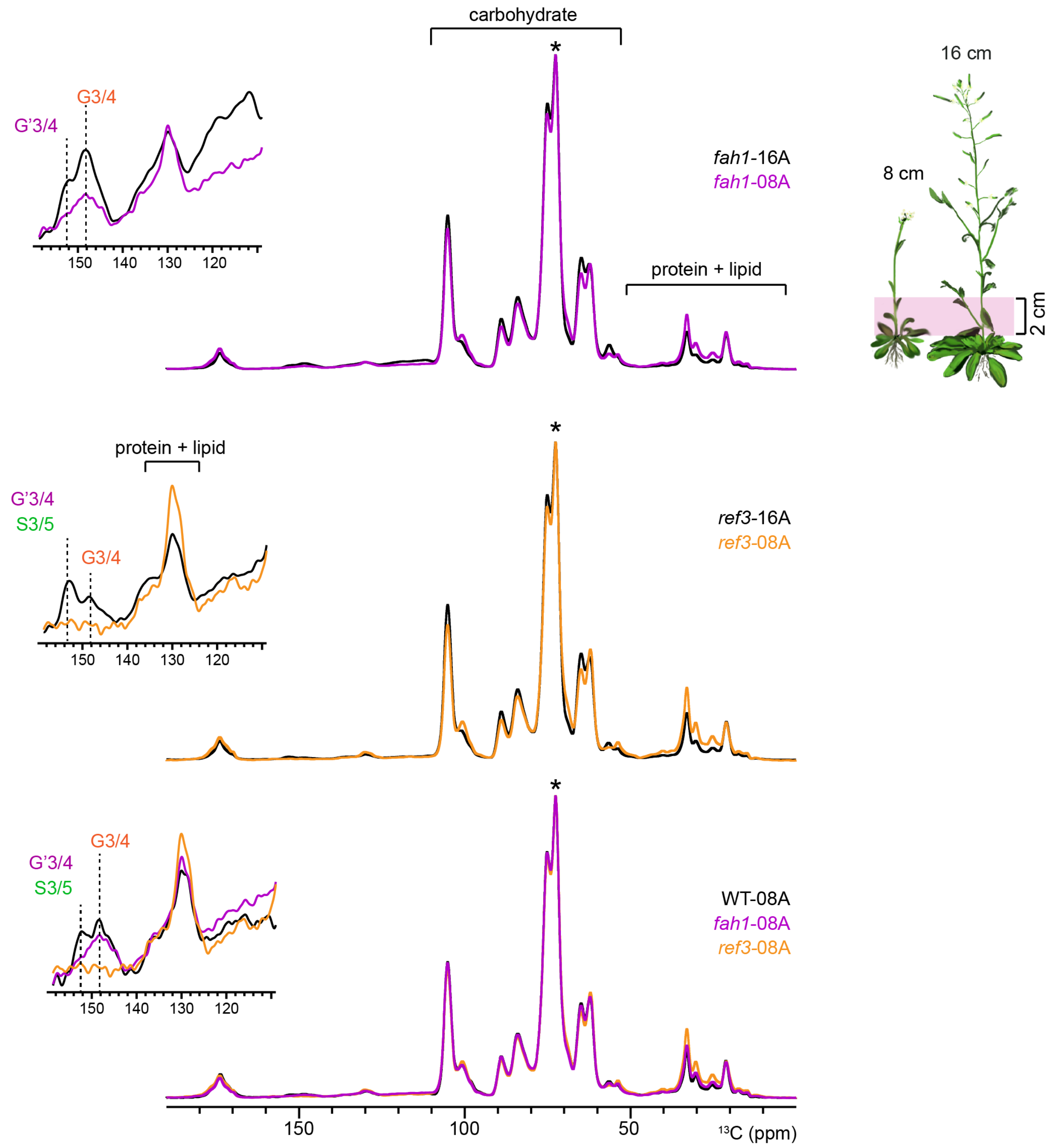


**Supplementary Figure 7. Comparison of lignin signals in mutants at different growth stages.** 1D CP spectra were collected on the basal segment A (0-2 cm) from inflorescences grown to heights of 8 cm and 16 cm, including the two mutants *fah1-2* and *ref3-3*, referred to as *fah1*-08/16A and *ref3*-08/16A, respectively. The 1D CP spectrum from WT 08A is also included in the overlay as a reference. All spectra are normalized with respect to the highest carbohydrate peak (asterisk). The zoomed-in region shows the difference in the lignin content, with the unique lignin peaks indicated by dashed lines. The unique lignin peaks are depleted in the *ref3*-08A sample, and the peak at around 130 ppm is dominated by aromatic residue and lipid signals. The protein and lipid signals in the 10-50 ppm region are also more prominent for the *ref3*-08A sample.

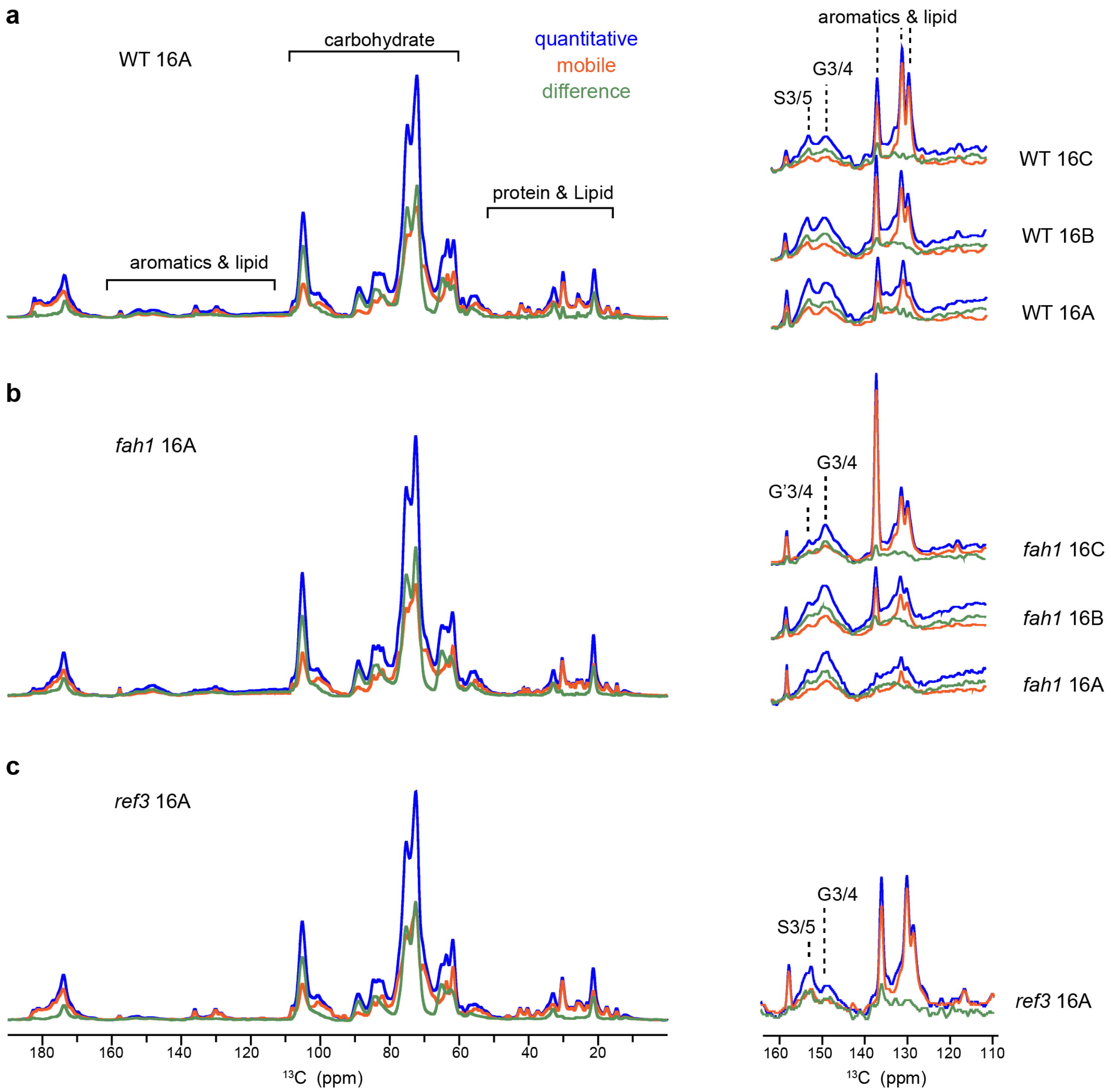


**Supplementary Figure 8. Quantitative DP spectra for WT and mutants of *Arabidopsis*.** 1D $^{13}$C DP spectra are overlaid for different segments of (**a**) WT, (**b**) *fah1* and (**c**) *ref3*. The spectra in blue have a long recycle delay of 35 s showing quantitative $^{13}$C composition. The spectra in red have a recycle delay of 2 s showing $^{13}$C species that are relatively more mobile and do not show up in the 1D CP spectra. The spectra in green show the difference between the two types of DP spectra, indicating relatively rigid fractions. The panels on the right display zoomed-in view of the region that include lignin aromatic carbons, fatty acyl chain carbons in lipids, and sidechain carbons of aromatic amino acid residues. The blue spectra on the right were normalized with respect to the highest carbohydrate peak at 72.5 ppm (not shown here) for each sample, and the red and green spectra were scaled according to their respective normalization factors. The strong sharp peaks at 136, 130 and 128.5 ppm in the zoomed-in region arise from the protein and lipid signals which are largely dynamic, and they grow stronger in higher segments from segment A to C. The lignin peaks are at around 152.5 ppm (S3/5, G'3/4) and 148.5 ppm (G3/4) and are mostly in the rigid fraction (green), apart from the *fah1*-16C and *ref3*-16A samples.

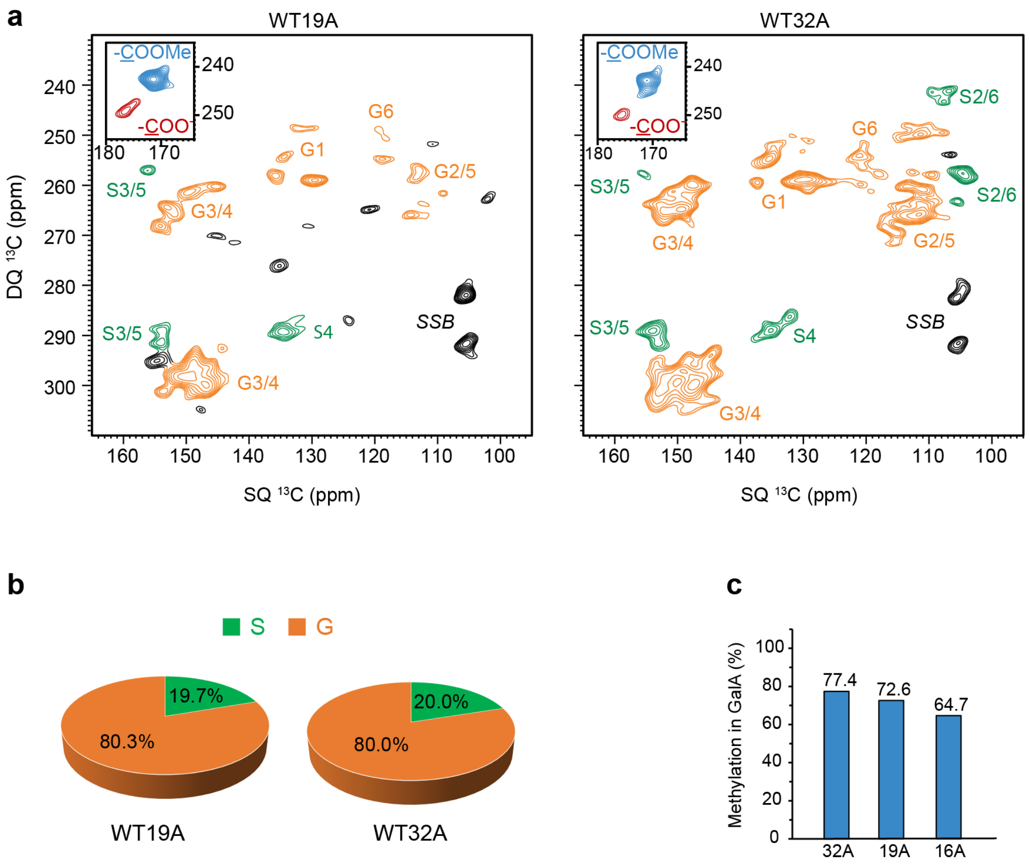


**Supplementary Figure 9. Additional CP J-INADEQUATE spectra of wild-type stems.** (**a**) CP-based J-INADEQUATE spectra collected on WT-19A (left) and WT-32A (right). The S and G lignin peaks are colored in green and orange, respectively. The inset shows the C6 peak region of GalA, with unmethylated in red and methylated in blue. The peaks in region 100 – 110 ppm (SQ)/ 280 – 295 ppm (DQ) are spinning sideband (SSB). (**b**) The S/G unit ratio in each sample. Estimations were based on the peak volumes corresponding to carbon 3/5 and 4 of the S units, and carbon 3/4 signals of the G units. (**c**) Methylation level of GalA in each sample. The molar fraction of methylated GalA was estimated by comparing the peak volumes of two carbonyl peaks corresponding to methylated and unmethylated GalA units shown in the inset of panel (a). The WT-16A data was also included here as a reference.

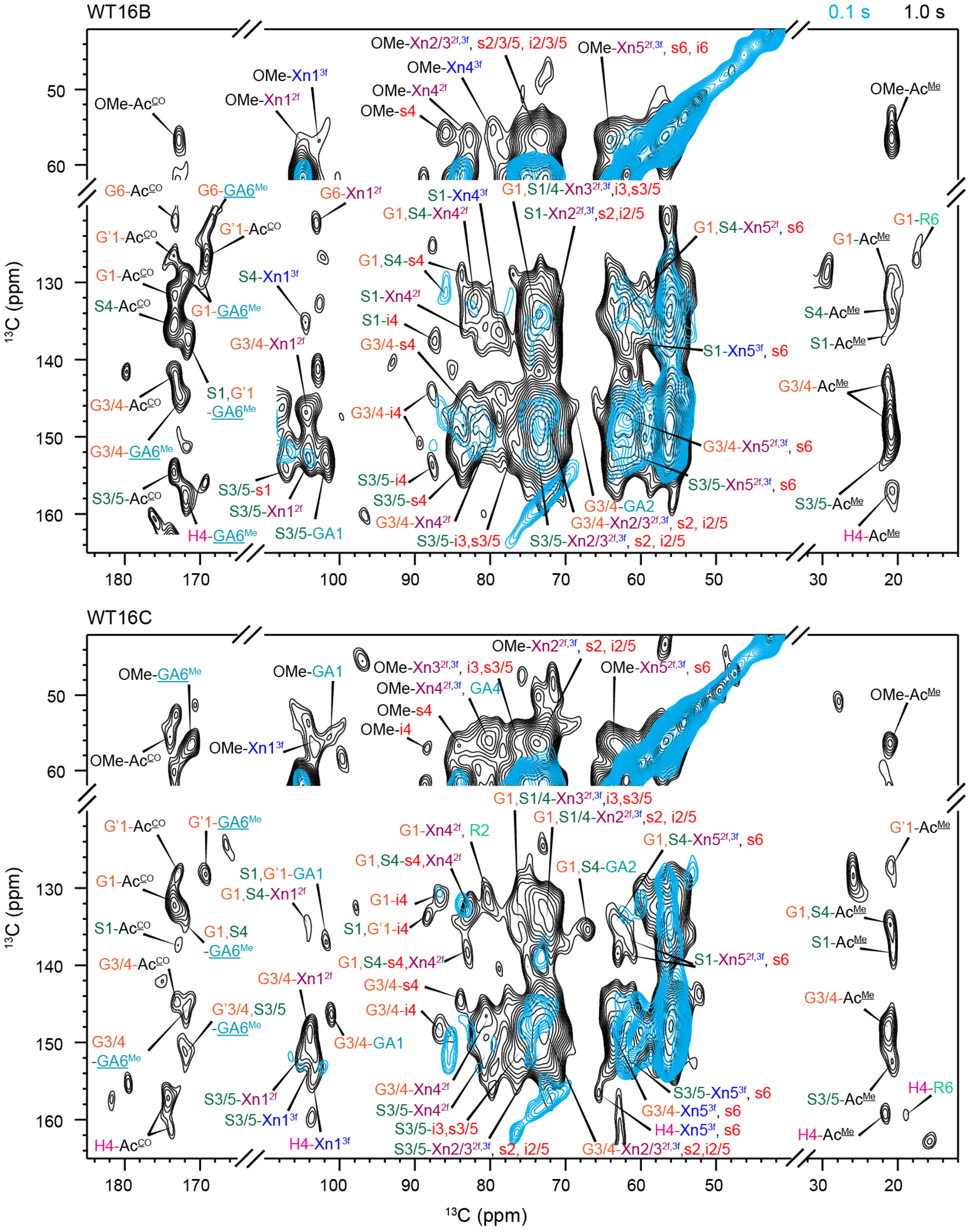


**Supplementary Figure 10. 2D $^{13}$C-$^{13}$C correlation spectra of wild-type samples.** For each sample, the spectra were measured using short (0.1 s; blue) and long (1.0 s; black) mixing times. Data are shown for WT 16B (top) and WT 16C (bottom) samples. Only lignin-carbohydrate cross-peak cross peaks are labeled. In addition to lignin-xylan and lignin-cellulose interactions, interactions between lignin and pectin can also be observed, for example, for the carbon sites of methylated GalA (GA6$^{Me}$, 168-172 ppm) and unmethylated GalA (GA6, 175.5-177.5 ppm), C1 and C2 of GalA (GA1 at 99.5-101.5 ppm and GA2 at 67-69 ppm), and C6 of rhamnose (R6, 17-19 ppm). Potential C2 peak of rhamnose (R2, 80 ppm) can also be observed but it is overlapping with Xn4$^{2f}$ (81.5 ppm) and Xn4$^{3f}$ (78.5 ppm).

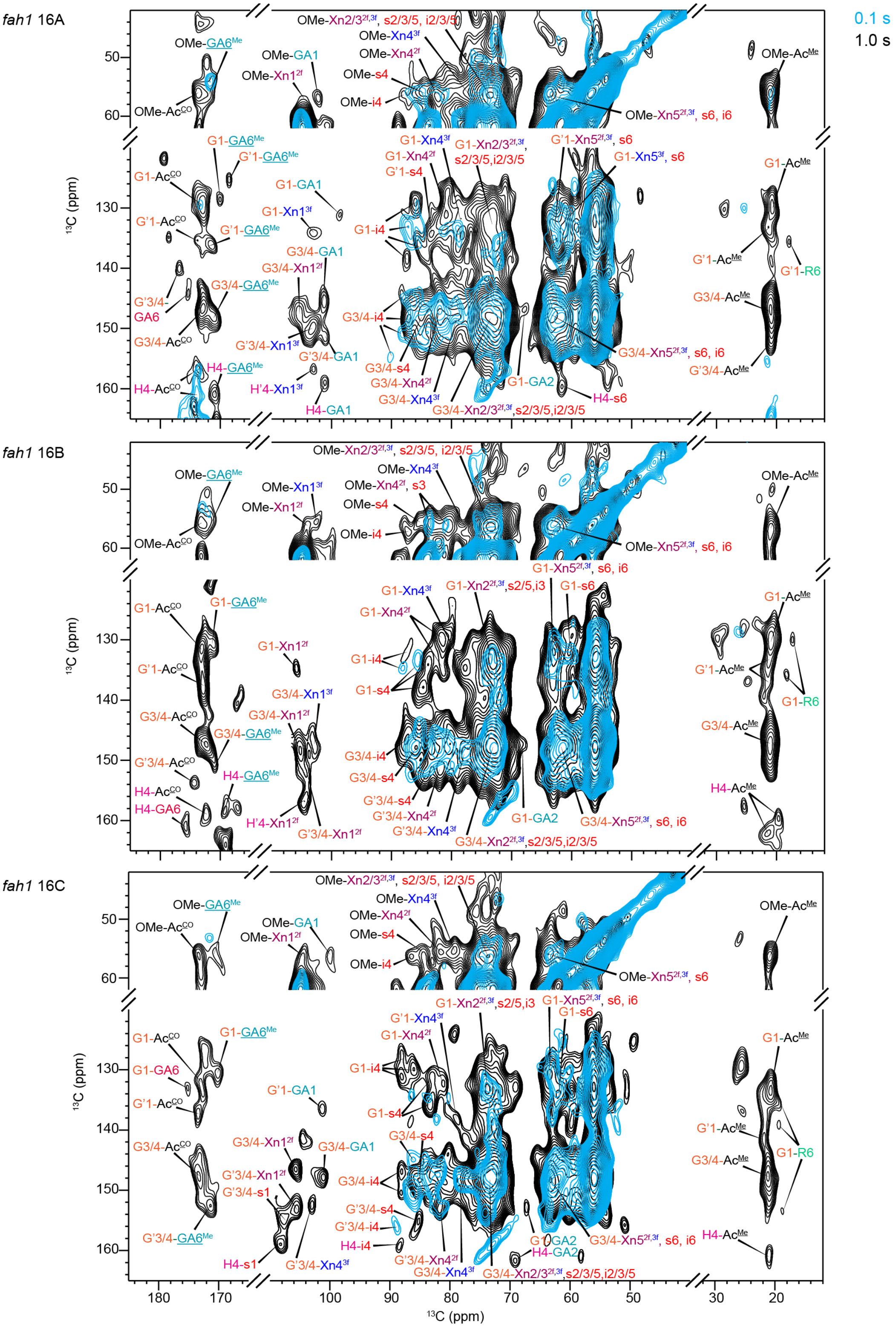


**Supplementary Figure 11. 2D $^{13}C$-$^{13}C$ correlation spectra of *fah1* samples.** For each sample, the spectra were measured using short (0.1 s; blue) and long (1.0 s; black) mixing times. Data are shown for *fah1*-16A (top), *fah1*-16B (middle), and *fah1*-16C (bottom). Lignin-carbohydrate cross-peak cross peaks are labeled.

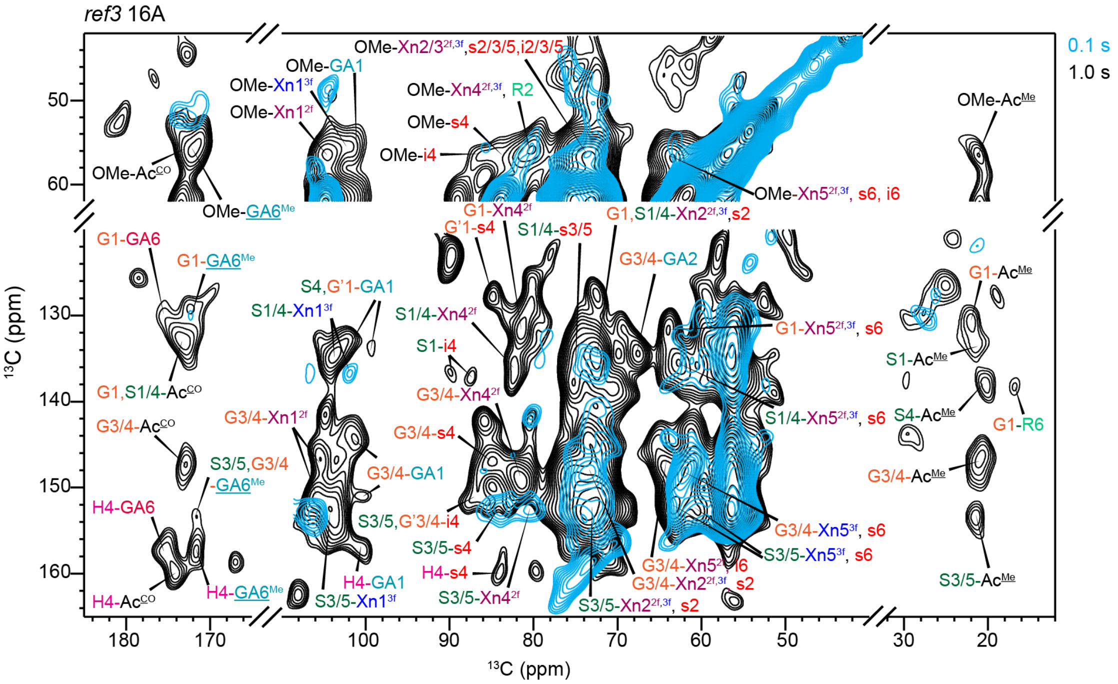


**Supplementary Figure 12. 2D $^{13}C$-$^{13}C$ correlation spectra of *ref3*-16A sample.** The spectra were measured using short (0.1 s; blue) and long (1.0 s; black) mixing times. Only lignin-carbohydrate cross-peak cross peaks are labeled.

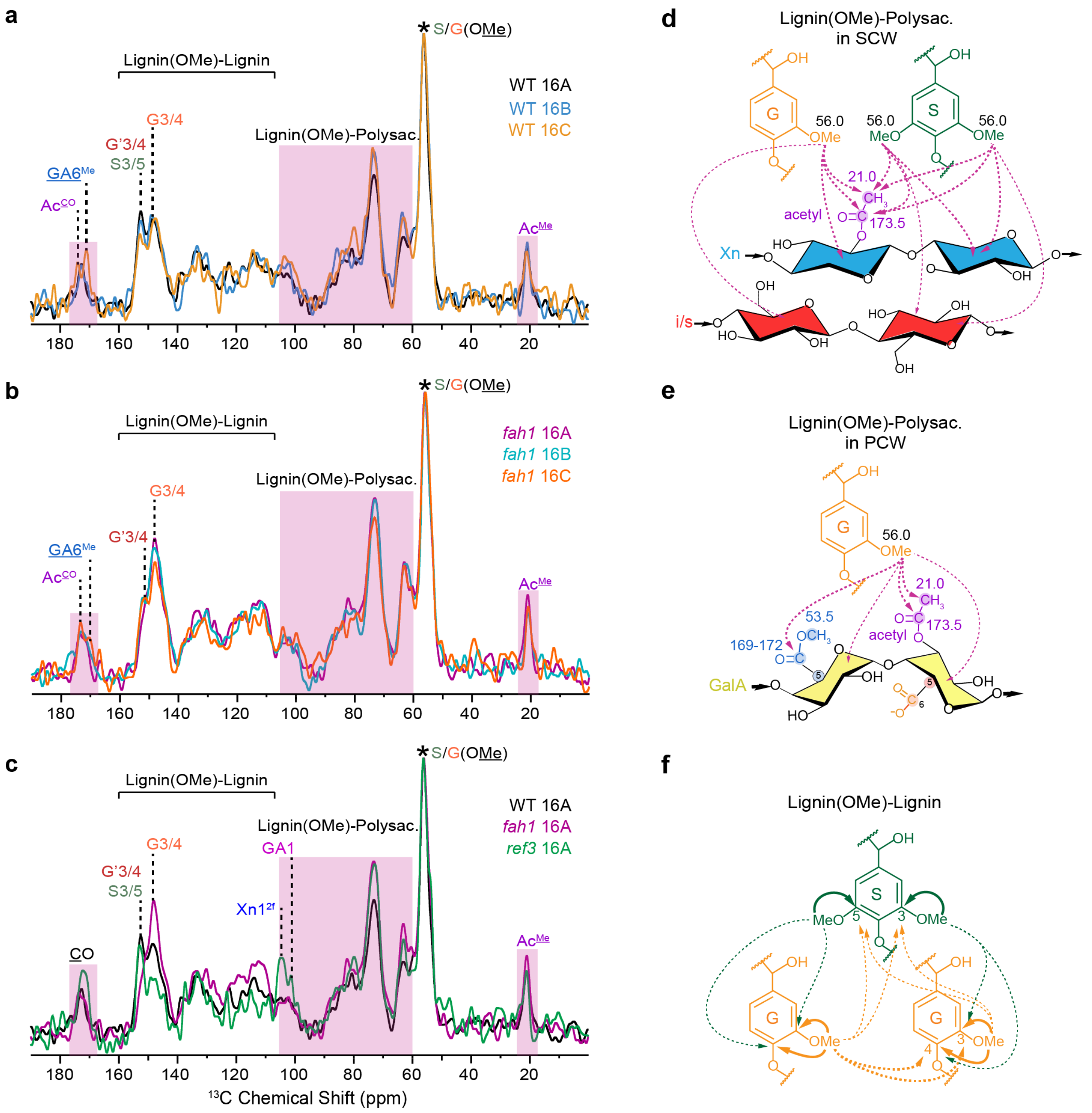


**Supplementary Figure 13. Intermolecular interactions viewed from 1D cross sections. (a-c)** 1D cross-sections extracted at 56 ppm (OMe) from 2D dipolar-gated PDSD spectra normalized by the OMe peak (asterisk). Lignin-carbohydrate interaction regions are highlighted in pink. Various carbonyl sites, i.e., acetyl groups ($Ac^{CO}$, 173.5 ppm) and methylated GalA C6 ($GA6^{Me}$, 171 ppm), are observed in the basal segment C of WT and *fah1*. Strong interactions of lignin with xylan ($Xn1^{2f}$, 104.5 ppm) and with pectin (GA1, 100.5 ppm) are observed in *ref3*-16A. (**d-f**) Structural representation of interactions observed in 1D slices. Inter- and intramolecular interactions are indicated by dashed lines and solid lines with arrows, respectively. (**d**) Interactions between lignin OMe groups and cellulose/xylan carbons. (**e**) Interactions between lignin OMe groups and pectin, including C1-C5 of GalA (GA), acetyl carbons of GalA, and methyl esters ($GA6^{Me}$, 171 ppm) from methylated GalA. (**f**) Interactions between lignin OMe groups and lignin aromatic carbon sites. Solid lines indicate the nearest intramolecular interactions, giving rise to the most prominent lignin aromatic carbon peaks in the 1D slices, G3/4 and S3/5. Signals from neighboring lignin units and those further away contribute to these peaks to a lesser extent and are indicated by dashed lines.

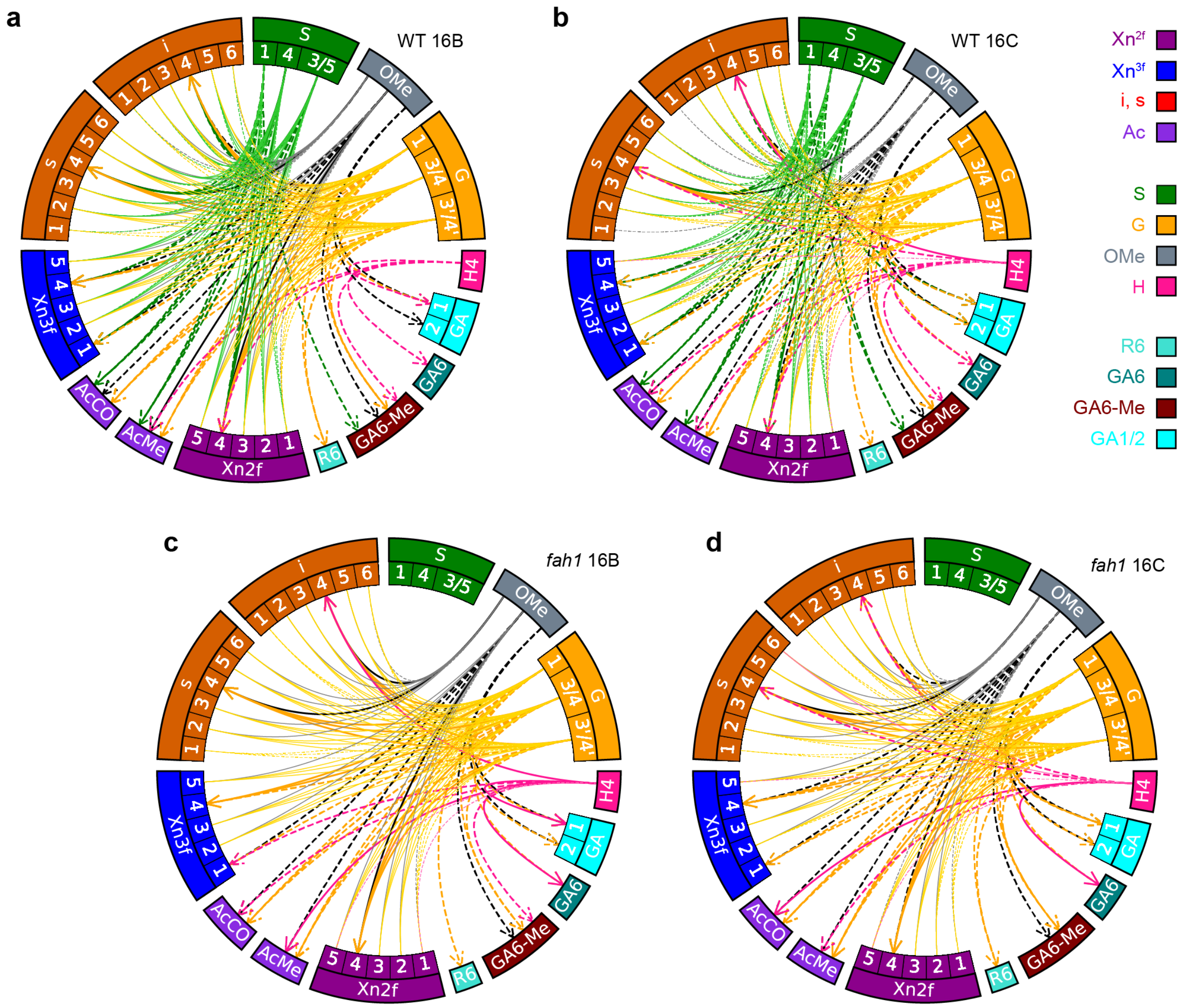


**Supplementary Figure 14. Chord diagram for intermolecular interactions.** The interactions were based on cross-peaks identified in the 2D dipolar-gated $^{13}C$-$^{13}C$ correlation spectra for samples (**a**) WT16B, (**b**) WT16C, (**c**) *fah1*-16B, and (**d**) *fah1*-16C. The carbon sites are color-coded with the legends on the right. Solid lines and dashed lines represent cross-peaks identified in the short-range and long-range spectra, respectively. Thick lines with arrow indicate unambiguous, site-specific cross-links and narrow lines without arrow indicate convoluted cross-links. The diagrams were compiled using pyCirclize script (https://github.com/moshi4/pyCirclize).

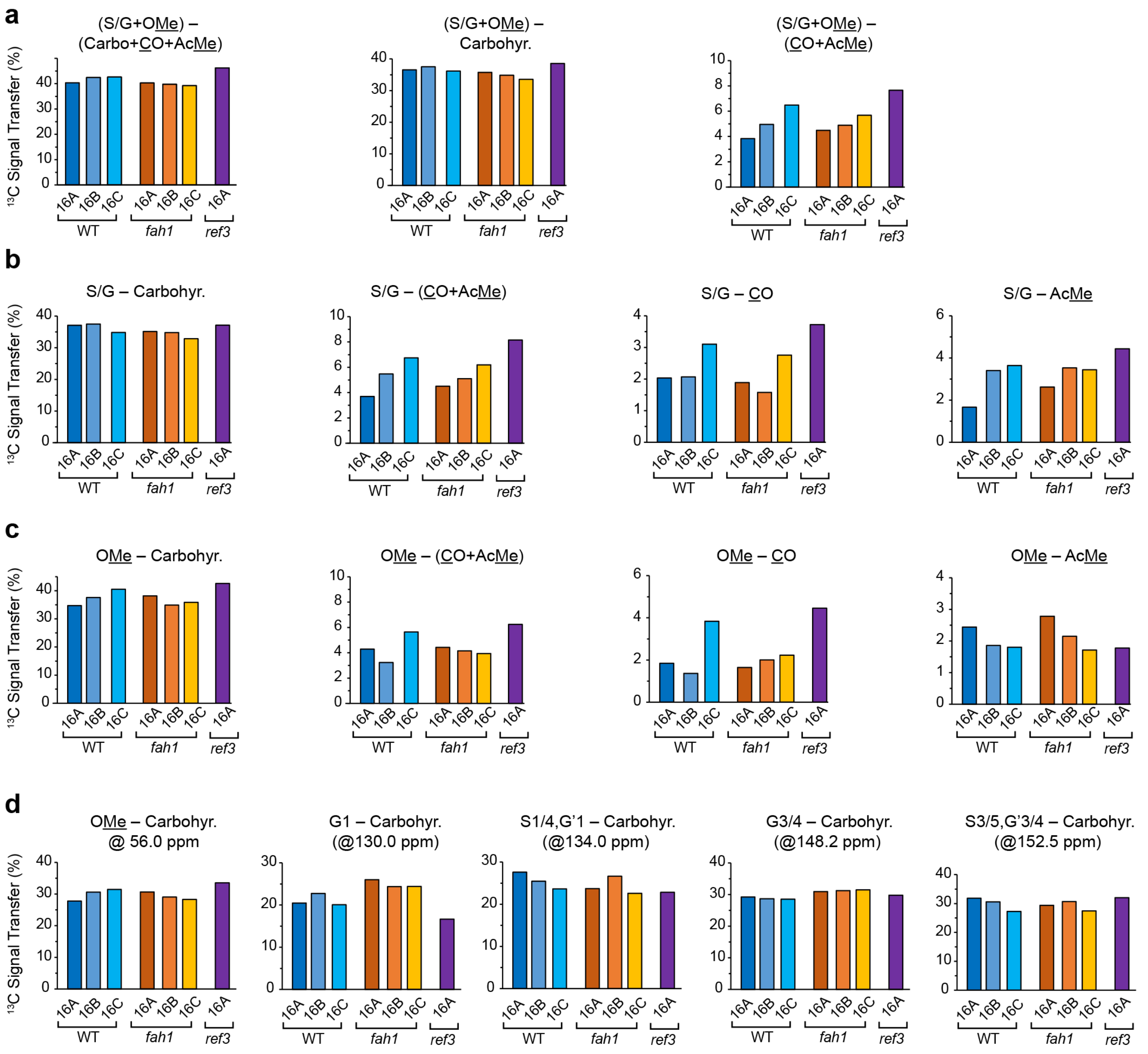


**Supplementary Figure 15. Lignin–carbohydrate interactions presented as bar charts.** Bar charts show the percentage carbon signals transferred from lignin to carbohydrate, analyzed from dipolar-gated PDSD spectra (1 s mixing time, long-range). The lignin-carbohydrate interactions were analyzed in four different categories: the overall lignin-carbohydrate contacts that include both S/G and OMe carbons (**a**), the transfer that only from S/G carbons (**b**) and OMe carbons (**c**), respectively, and the transfer from specific carbon sites analyzed from 1D cross-sections (**d**). S/G: deprotonated lignin carbons; OMe: lignin methoxy carbons; carbo: carbohydrate carbons with acetyls excluded; Ac: acetyl methyl and carbonyl carbons; AcMe: methyl carbons from acetyl group; CO: carbonyl carbons, including those from acetyl and C6 of GalA. The integration regions for each group are summarized in **Supplementary Table 5**. The resolved lignin carbon sites and their chemical shift values where 1D cross-sections were extracted for the analysis are indicated on top of each chart in panel (**d**).

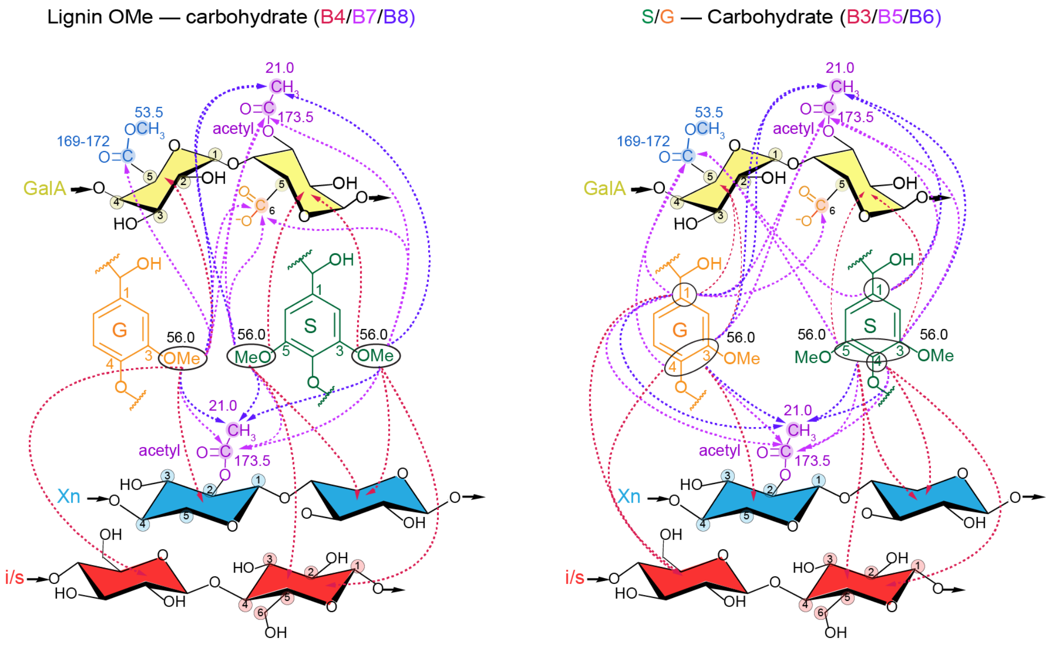


**Supplementary Figure 16. Key interactions of lignin OMe and aromatics with carbohydrates.** The interactions between lignin OMe and carbohydrates (cellulose: red; xylan: blue; GalA: yellow) are illustrated on the left, corresponding to spectral regions of B4, B7 and B8 listed in **Supplementary Table 5**; the interactions between deprotonated lignin carbons and carbohydrates are on the right, corresponding to spectral regions of B3, B5 and B6.

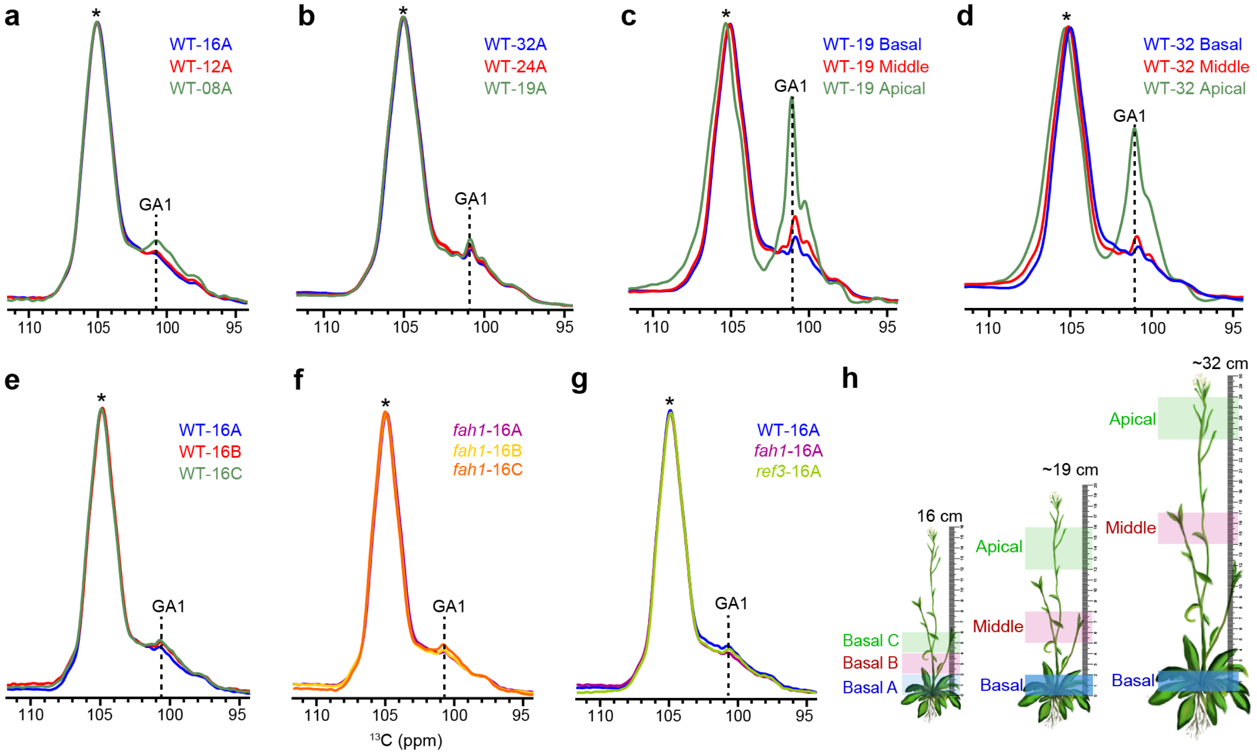


**Supplementary Figure 17. Comparison of the rigid pectin signal in all samples.** 1D CP spectra are overlayed to show changes of pectin signal in the rigid fraction. Only the C1 peak region of polysaccharides are displayed, in which the pectin signals can be resolved and distinguished from other carbohydrates. All spectra are normalized with respect to the C1 peak of xylan and cellulose (Xn/i/s, asterisk) at around 105 ppm. The C1 peaks of GalA (GA1) at around 100.5 ppm are indicated by dashed lines. (**a**) Comparison between samples WT-16A/12A/08A. (**b**) Comparison between samples WT-32A/24A/19A. (**c**) Comparison between different segments of the WT 19 cm sample. (**d**) Comparison between different segments of the WT 32 cm sample. (**e**) Comparison between samples WT-16A/B/C. (**f**) Comparison between samples *fah1*-16A/B/C. (**g**) Comparison of samples WT16A/*fah1*-16A/*ref3*-16A. (**h**) Illustration of the segments used for each sample.

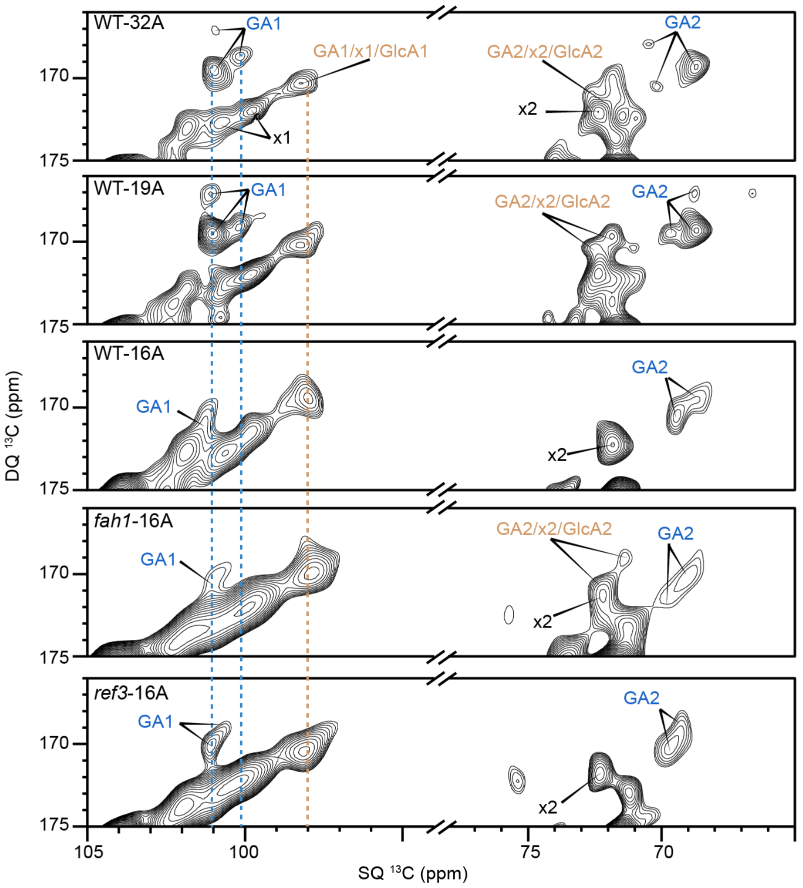


**Supplementary Figure 18. Identification of GalA and GlcA signals.** C1-C2 region of CP-based refocused J-INADEQUATE spectra showing signals of GalA (GA) and GlcA for samples of WT-32A, WT-19A, WT-16A, *fah1*-16A and *ref3*-16A. The main GA1 SQ chemical shifts (101.5-99.5 ppm) are indicated by the blue dashed lines, which partially overlap with that of xylose (x1) units in xyloglucan. The GlcA1 SQ chemical shift (98 ppm), indicated by the brown dashed line, potentially overlaps with minor species of GA1 and x1.

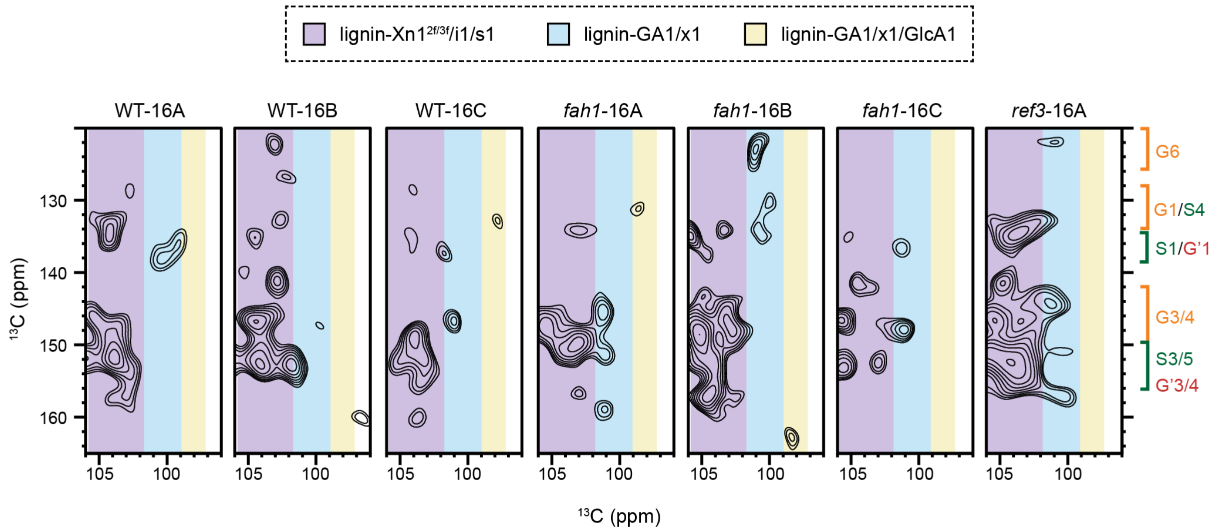


**Supplementary Figure 19. Absence of lignin-GlcA cross peaks.** Spectral regions of gated long-range $^{13}$C-$^{13}$C correlation spectra show interactions between lignin aromatic carbons and various C1 sites. The purple band indicates lignin interactions with xylan and cellulose. The blue band highlights lignin interactions with pectin, with potential mixed contributions from xyloglucan. The yellow band highlights lignin interactions with GlcA, with potential contributions from minor species of GA1 and x1, where no prominent interactions are observed.

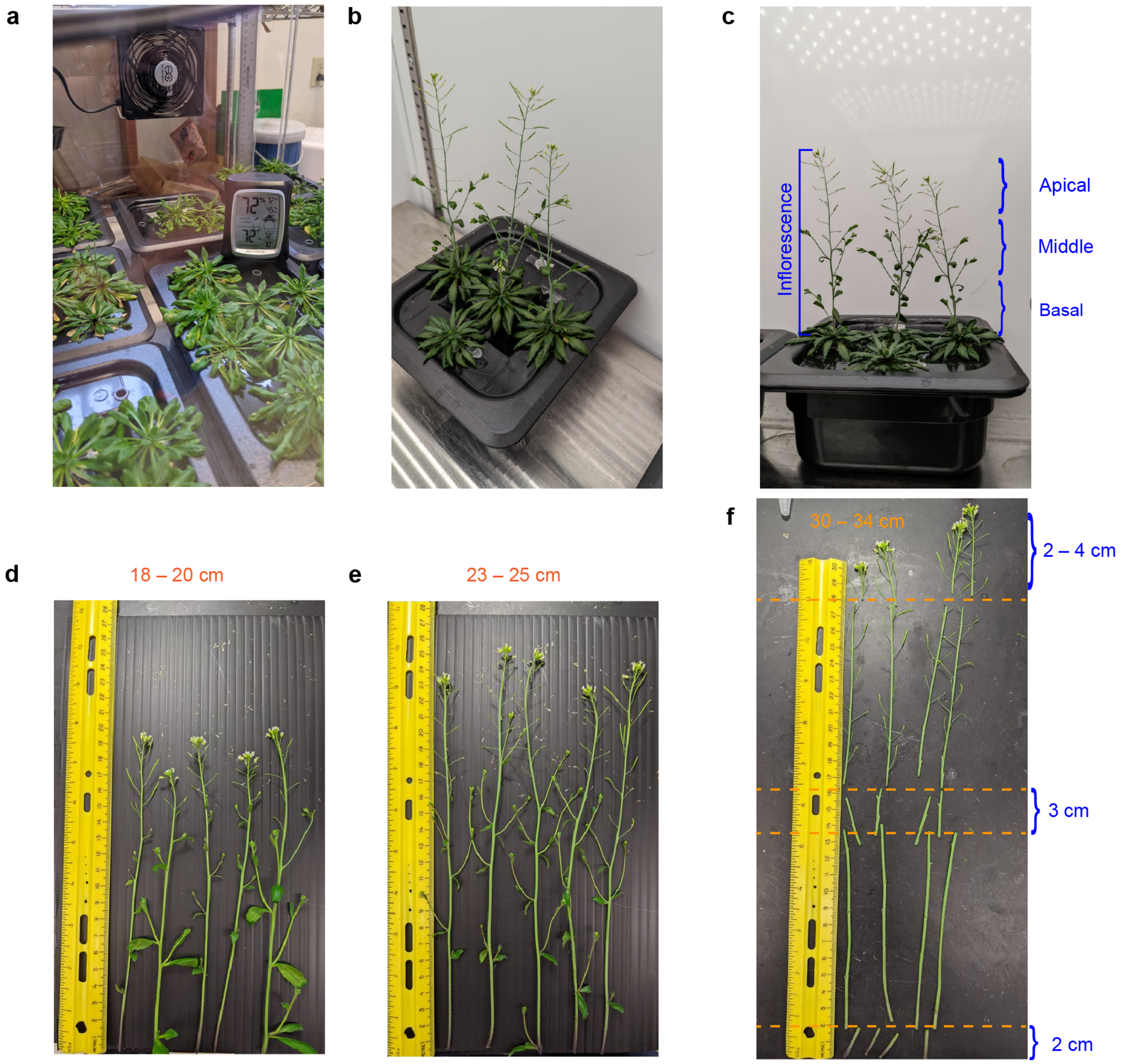


**Supplementary Figure 20. Illustrations of *Arabidopsis* stem sample preparation.** (**a**) Hydroponic Growth of *Arabidopsis*. (**b-c**) Top view and side view of *Arabidopsis* inflorescence, respectively. The basal, middle and apical regions are indicated on the right. (**d-f**) Photographs of *Arabidopsis* stems harvested at different growth stages, with the lengths of the stems indicated at the top. (**f**) shows an example of how different regions of stems were cut for preparing the samples used in the NMR experiments.

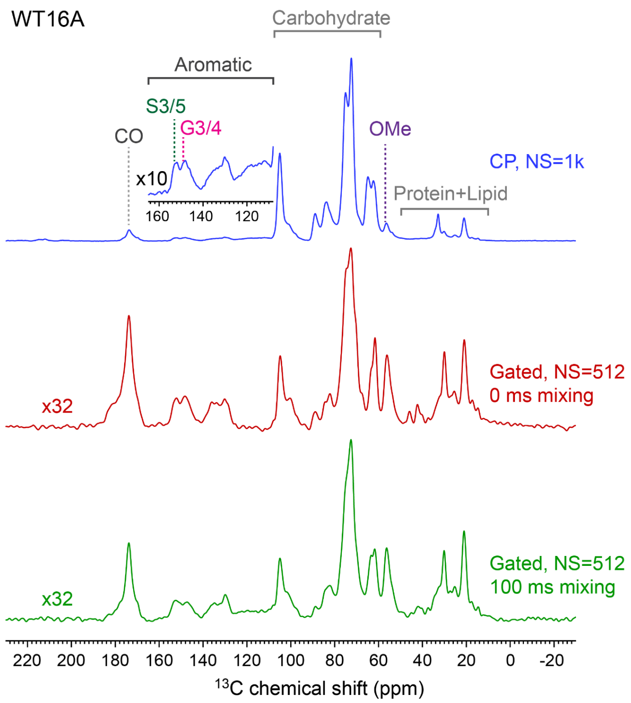


**Supplementary Figure 21. 1D spectra showing the effect of dipolar gating.** Top row is the 1D CP experiment without the dipolar-gating step, in which the carbohydrate region is about a hundredfold more intense than the aromatic region; the latter is barely visible without the 10 times magnification in the inset. The middle and bottom rows are the 1D spectra with incorporation of the dipolar-gating step, both of which are scaled up by a factor of 32. The intensity ratio between the carbohydrate region and the aromatic region is attenuated by tenfold in both dipolar-gated spectra.

**Supplementary Table 1. Number of intermolecular cross peaks between carbohydrates and lignin.**

| | OMe-<br>i/s | OMe -<br>Xn2f | OMe -<br>Xn3f | OMe -<br>Ac(CO+Me) | OMe -<br>pectin | Total #<br>OMe -Carbohydrate |
|---|---|---|---|---|---|---|
| WT16A | 10 | 5 | 5 | 2 | 2 | 24 |
| WT16B | 10 | 5 | 5 | 2 | 3 | 25 |
| WT16C | 10 | 5 | 5 | 2 | 3 | 25 |
| *ref3*-16A | 10 | 5 | 5 | 2 | 3 | 25 |
| *fah1*-16A | 8 | 5 | 5 | 2 | 2 | 22 |
| *fah1*-16B | 8 | 5 | 4 | 2 | 3 | 22 |
| *fah1*-16C | 8 | 5 | 5 | 2 | 3 | 23 |
| | # S-<br>i/s | S-<br>Xn2f | S-<br>Xn3f | S-<br>Ac(CO+Me) | S-<br>pectin | Total #<br>S-Carbohydrate |
| WT16A | 23 | 14 | 13 | 6 | 4 | 60 |
| WT16B | 24 | 15 | 14 | 6 | 1 | 60 |
| WT16C | 19 | 14 | 13 | 5 | 4 | 55 |
| *ref3*-16A | 23 | 15 | 15 | 4 | 5 | 62 |
| *fah1*-16A | | | | | | |
| *fah1*-16B | | | | | | |
| *fah1*-16C | | | | | | |
| | # G-<br>i/s | G-<br>Xn2f | G-<br>Xn3f | G-<br>Ac(CO+Me) | G-<br>pectin | Total #<br>G-Carbohydrate |
| WT16A | 24 | 14 | 13 | 5 | 6 | 62 |
| WT16B | 27 | 15 | 12 | 6 | 6 | 66 |
| WT16C | 22 | 15 | 11 | 5 | 7 | 60 |
| *ref3*-16A | 23 | 15 | 12 | 5 | 7 | 62 |
| *fah1*-16A | 26 | 13 | 15 | 5 | 10 | 69 |
| *fah1*-16B | 30 | 15 | 13 | 6 | 4 | 68 |
| *fah1*-16C | 27 | 15 | 14 | 6 | 9 | 71 |
| | # H-<br>i/s | H-<br>Xn2f | H-<br>Xn3f | H-<br>Ac(CO+Me) | H-<br>pectin | Total #<br>H-Carbohydrate |
| WT16A | 2 | 2 | 1 | 1 | 3 | 9 |
| WT16B | 0 | 1 | 0 | 1 | 3 | 5 |
| WT16C | 2 | 2 | 1 | 2 | 3 | 10 |
| *ref3*-16A | 1 | 0 | 1 | 1 | 4 | 7 |
| *fah1*-16A | 2 | 1 | 3 | 1 | 4 | 11 |
| *fah1*-16B | 1 | 2 | 2 | 2 | 3 | 10 |
| *fah1*-16C | 3 | 1 | 0 | 2 | 2 | 8 |

**Supplementary Table 2. The *Arabidopsis* samples for NMR experiments.** The short name for each sample is indicated in the bracket after the cut range. When the full height of the inflorescence varies, the average value is given with the range of the length indicated in the bracket.

| **WT** | | | | | | |
|---|---|---|---|---|---|---|
| **Height (cm)** | **# of stems** | **Basal Cut Range (cm)** | | | **Middle Cut Range (cm)** | **Apical Cut Range (cm)** |
| 8 | 4 | 0 – 2 (08A) | | | | |
| 12 | 4 | 0 – 2 (12A) | | | | |
| 16 | 3 | 0 – 2 (16A) | 2 – 4 (16B) | 4 – 6 (16C) | | |
| 19 (18 – 20) | 5 | 0 – 2 (19A) | | | 8 – 11 (19Middle) | 16 – 18/20 (19Apical) |
| 24 (23 – 25) | 5 | 0 – 2 (24A) | | | | |
| 32 (30 – 34) | 4 | 0 – 2 (32A) | | | 14 – 17 (19Middle) | 28 – 30/30–34 (32Apical) |
| ***fah1-2*** | | | | | | |
| **Height (cm)** | **# of stems** | **Basal Cut Range (cm)** | | | - | - |
| 8 | 5 | 0 – 2 (08A) | | | - | - |
| 16 | 2 | 0 – 2 (16A) | 2 – 4 (16B) | 4 – 6 (16C) | - | - |
| ***ref3-3*** | | | | | | |
| **Height (cm)** | **# of stems** | **Basal Cut Range (cm)** | | | - | - |
| 8 | 5 | 0 – 2 (08A) | | | - | - |
| 16 | 4 | 0 – 2 (16A) | - | - | - | - |

**Supplementary Table 3. NMR experimental parameters for all samples.** T = probe temperature; $B_0$ = magnetic field; $\nu_r$ = MAS frequency; NS = number of scans; $d_1$ = recycle delay; $\tau_{HC}$ = initial $^1H/^{13}C$ CP contact time; $t_2/t_1$ = acquisition length of direct/indirect dimension; TD2/TD1 = total points of FID for the direct/indirect dimension acquisition; TDeff = effective FID data points used for Fourier transform; LB = line broadening parameter; GB = position of the maximum of the Gaussian function. The 2D acquisition mode for all 2D experiments was States. SPINAL64 $^1H$ decoupling was used for all experiments at a field strength of 83.3 kHz.

| Sample | Experiment | T (K) | $B_0$ (T) | $\nu_r$ (kHz) | NS | $d_1$ (s) | $\tau_{HC}$ (ms) | $t_2/t_1$ (ms) | TD2/TD1 | TDeff | LB (Hz) | GB |
|---|---|---|---|---|---|---|---|---|---|---|---|---|
| WT08A | 1D CP | 290 | 9.4 | 14 | 1024 | 2.0 | 1.0 | 16.0 | 1600 | - | -40 | 0.05 |
| | 2D Gated DARR | 290 | 9.4 | 14 | 480 | 1.5 | 1.0 | 12.0/2.9 | 1200/112 | 500/112 | -50/-50 | 0.05/0.05 |
| WT12A | 2D Gated PDSD | 290 | 9.4 | 14 | 320*2 | 1.1 | 1.0 | 10.2/2.1 | 1024/80 | 350/80 | -60/-60 | 0.05/0.05 |
| WT16A | 1D CP | 298 | 14.1 | 14 | 1024 | 2.0 | 1.0 | 16.0 | 1600 | - | -40 | 0.05 |
| | 1D DP | 298 | 14.1 | 14 | 256 | 35.0, 2.0 | - | 16.0 | 1600 | - | -10 | 0.05 |
| | 2D Gated DARR | 298 | 14.1 | 14 | 320 | 1.5 | 1.0 | 12./3.0 | 1200/162 | 600/162 | -60/-60 | 0.05/0.05 |
| | 2D Gated PDSD | 298 | 14.1 | 14 | 512 | 1.1 | 1.0 | 10.2/2.1 | 1024/116 | 600/116 | -60/-60 | 0.05/0.05 |
| | 2D CP J-INADEQUATE | 275 | 14.1 | 18 | 360*2 | 1.5 | 1.9 | 10.0/3.2 | 1000/250 | 300/250 | -50/-50 | 0.05/0.05 |
| WT16B | 1D CP | 298 | 14.1 | 14 | 1024 | 2.0 | 1.0 | 16.0 | 1600 | - | -40 | 0.05 |
| | 1D DP | 298 | 14.1 | 14 | 256 | 35.0, 2.0 | - | 16.0 | 1600 | - | -10 | 0.05 |
| WT16C | 2D Gated DARR | 298 | 14.1 | 14 | 384 | 1.5 | 1.0 | 12./3.0 | 1200/162 | 600/162 | -60/-60 | 0.05/0.05 |
| | 2D Gated PDSD | 298 | 14.1 | 14 | 512 | 1.1 | 1.0 | 10.2/2.1 | 1024/116 | 600/116 | -80/-60 | 0.03/0.05 |
| WT19A | 1D CP | 293 | 16.4 | 20 | 4096 | 1.8 | 1.0 | 10.0 | 1250 | - | -10 | 0.05 |
| | 1D DP | 293 | 16.4 | 20 | 512 | 35.0 | - | 16.4 | 2048 | - | -10 | 0.05 |
| | 2D CP J-INADEQUATE | 293 | 16.4 | 20 | 64*2 | 1.8 | 1.0 | 12.8/5.2 | 1600/400 | 400/270 | -50/-50 | 0.05/0.05 |
| WT19 Middle | 1D CP | 293 | 16.4 | 20 | 4096 | 1.8 | 1.0 | 10.0 | 1250 | - | -10 | 0.05 |
| WT19 Apical | 1D DP | 293 | 16.4 | 20 | 1024 | 35.0 | - | 16.4 | 2048 | - | -10 | 0.05 |
| WT24A | 1D CP | 293 | 16.4 | 20 | 6144 | 1.8 | 1.0 | 9.7 | 1024 | - | -10 | 0.05 |
| | 1D DP | 293 | 16.4 | 20 | 1024 | 35.0 | - | 16.4 | 2048 | - | -10 | 0.05 |
| WT32A | 1D CP | 293 | 16.4 | 20 | 4096 | 1.8 | 1.0 | 10.0 | 1250 | - | -10 | 0.05 |
| | 1D DP | 293 | 16.4 | 20 | 1024 | 35.0 | - | 16.4 | 2048 | - | -10 | 0.05 |
| | 2D CP J-INADEQUATE | 293 | 16.4 | 20 | 64*2 | 1.8 | 1.0 | 12.8/5.2 | 1600/400 | 400/270 | -50/-50 | 0.05/0.05 |
| WT32 Middle | 1D CP | 293 | 16.4 | 20 | 4096 | 1.8 | 1.0 | 10.0 | 1250 | - | -10 | 0.05 |
| WT32 Apical | 1D DP | 293 | 16.4 | 20 | 1024 | 35.0 | - | 16.4 | 2048 | - | -10 | 0.05 |
| *fah1* 08A | 1D CP | 290 | 9.4 | 14 | 1024 | 2.0 | 1.0 | 16 | 1600 | - | -10 | 0.05 |
| *fah1* 16A | 1D CP | 298 | 14.1 | 14 | 1024 | 2.0 | 1.0 | 16.0 | 1600 | - | -40 | 0.05 |
| | 1D DP | 298 | 14.1 | 14 | 256 | 35.0, 2.0 | - | 16.0 | 1600 | - | -10 | 0.05 |
| | 2D Gated DARR | 298 | 14.1 | 14 | 384 | 1.5 | 1.0 | 12./3.0 | 1200/162 | 600/162 | -60/-60 | 0.05/0.05 |
| | 2D Gated PDSD | 298 | 14.1 | 14 | 512 | 1.1 | 1.0 | 10.2/2.1 | 1024/116 | 600/116 | -80/-60 | 0.03/0.05 |
| | 2D CP J-INADEQUATE | 275 | 14.1 | 18 | 360*2 | 1.5 | 2.1 | 10.0/3.2 | 1000/250 | 400/250 | -50/-50 | 0.05/0.05 |
| | 1D CP | 298 | 14.1 | 14 | 1024 | 2.0 | 1.0 | 16.0 | 1600 | - | -40 | 0.05 |
| *fah1* 16B | 1D DP | 298 | 14.1 | 14 | 256 | 35.0, 2.0 | - | 16.0 | 1600 | - | -10 | 0.05 |
| *fah1* 16C | 2D Gated DARR | 298 | 14.1 | 14 | 384 | 1.5 | 1.0 | 12./3.0 | 1200/162 | 600/162 | -60/-60 | 0.05/0.05 |
| | 2D Gated PDSD | 298 | 14.1 | 14 | 512 | 1.1 | 1.0 | 10.2/2.1 | 1024/116 | 600/116 | -80/-60 | 0.03/0.05 |
| *ref3* 08A | 1D CP | 290 | 9.4 | 14 | 1024 | 2.0 | 1.0 | 16 | 1600 | - | -10 | 0.05 |
| *ref3* 16A | 1D CP | 298 | 14.1 | 14 | 1024 | 2.0 | 1.0 | 16.0 | 1600 | - | -40 | 0.05 |
| | 1D DP | 298 | 14.1 | 14 | 256 | 35.0, 2.0 | - | 16.0 | 1600 | - | -10 | 0.05 |
| | 2D Gated DARR | 298 | 14.1 | 14 | 384 | 1.5 | 1.0 | 12./3.0 | 1200/162 | 800/162 | -60/-60 | 0.05/0.05 |
| | 2D Gated PDSD | 298 | 14.1 | 14 | 512 | 1.1 | 1.0 | 10.2/2.1 | 1024/116 | 400/116 | -80/-80 | 0.03/0.03 |
| | 2D CP J-INADEQUATE | 275 | 14.1 | 18 | 360*2 | 1.5 | 2.0 | 10.0/3.2 | 1000/250 | 400/250 | -50/-50 | 0.05/0.05 |

**Supplementary Table 4. $^{13}$C chemical shifts of lignin on TMS scale.** Unidentified site is indicated as "-". Degenerate sites are separated by "/".

| | G | | | | | | S | | | | | |
|---|---|---|---|---|---|---|---|---|---|---|---|---|
| Sample | C1 | C2 | C3* | C4* | C5 | C6 | C1 | C2 | C3 | C4 | C5 | C6 |
| WT-16A | 134.8 | 111.2 | 147.6 | 144.0 | 110.9 | 118.8 | 138.3 | 105.5 | 152.8 | 134.5 | 152.8 | 105.5 |
| | 134.8/136.9 | 112.0 | 148.7 | 145.6 | 115.5/117.2 | - | 135.5 | 104.3 | 153.9 | 133.1 | 153.9 | 104.3 |
| | 133.4 | 111.4/109.5 | 152.4 | 146.6 | 115.5/117.2 | - | | | | | | |
| | 130.5 | 114.2 | 151.1 | 148.5 | 116.5 | - | | | | | | |
| WT-19A | 131.8 | 112.5 | 148.5 | 144.6 | 115.6 | - | 135.3 | 108.6 | 153.5 | 135.2 | 153.5 | 108.6 |
| | 131.8 | 112.6 | 149.2 | 145.8 | 114.1 | 121.9 | 135.3 | 108.3 | 154.1 | 137.3 | 154.1 | 108.3 |
| | 138.9 | 111.5 | 149.9 | 147.4 | 118.5 | - | 135.1 | 104.2 | 154.7 | 131.1 | 154.7 | 104.2 |
| | 132.5 | 114.5/115.7 | 150.9 | 148.5 | 112.7 | 118.8/122.5 | 135.0/138.1 | 106.6 | 155.0 | 134.2 | 155.6 | 106.6 |
| | 133.6 | 113.7 | 151.9 | 146.5 | 116.2 | - | | | | | | |
| | 133.6 | 113.5 | 153.8 | 147.2 | 117.5 | - | | | | | | |
| WT-32A | 130.3/136.6 | 111.6/109.5 | 148.4 | 145.1 | - | - | - | 104.6/108.5 | 152.9 | 135.4 | 152.9 | 104.6/108.5 |
| | 136.4 | 114.6 | 149.4 | 146.1 | 113.8 | - | - | 103.8 | 154.7 | 134.4 | 154.7 | 103.8 |
| | 136.4 | 113.9 | 152.0 | 146.4 | 113.8 | - | - | 104.6/108.5 | 153.2 | 137.6 | 153.2 | 104.6/108.5 |
| | 136.3/135.4 | 110.7/116.8 | 151.3 | 148.7 | 116.5 | - | - | 104.6/108.5 | 153.7 | 132.2 | 154.0 | 104.6/108.5 |
| | 132.7/134.7 | 112.8/110.8 | 152.8 | 147.0 | 116.7/113.5 | - | - | 104.6/108.5 | 153.3 | 141.5 | 153.3 | 104.6/108.5 |
| | 134.7 | - | - | - | - | 119.8 | - | - | 153.6 | 144.1 | 153.6 | - |
| | 132.6 | - | - | - | - | 120.5 | 134.8 | 106.5 | - | - | - | 106.5 |
| | - | - | 153.6 | 144.1 | - | - | | | | | | |
| | - | - | 153.6 | 144.1 | - | - | | | | | | |
| *ref3-16A* | 135.7/132.1 | 113.2/111.5 | 147.7 | 143.5 | - | - | 136.4 | 103.3 | 153.2 | 134.0 | 153.2 | 103.3 |
| | 135.7/132.1 | 113.2/111.5 | 148.7 | 144.6 | 118.9 | - | 136.4/134.5 | 103.5/108.6 | 153.8 | 132.9 | 153.8 | 103.5/108.6 |
| | 135.7/132.1 | 113.2/111.5 | 148.5 | 145.8 | 118.5 | - | 137.0 | 102.7 | 152.6 | 136.0 | 152.6 | 102.7 |
| | 133.6 | 113.5 | 150.8 | 146.4 | 117.8 | 124.2 | 137.0 | 102.8 | 152.4 | 138.8 | 152.4 | 102.8 |
| | 136.6 | 110.4 | 150.4 | 147.7 | 115.7 | - | - | - | 151.2 | 140.8 | 151.2 | - |
| | 129.2 | 112.4 | 153.0 | 146.5 | 117.8 | 124.2 | 136.2 | 106.2 | - | - | - | 106.2 |
| | 129.2 | 112.4 | 152.7 | 148.2 | 115.3 | - | | | | | | |
| | 129.2 | 112.4 | 152.3 | 149.6 | 116.9 | - | | | | | | |
| *fah1-16A* | 136.2/138.1 | 109.6/112.9 | 148.8 | 146.1 | - | - | | | | | | |
| | 136.8 | 113.8/114.8 | 149.7 | 146.3 | - | - | | | | | | |
| | 129.8/136.2 | 113.6/116.7 | 150.1 | 147.6 | 114.0 | 119.5 | | | | | | |
| | 133.1/136.6 | 112.8/114.8 | 152.1 | 146.8 | 115.5 | - | | | | | | |
| | 133.1/138.1 | 112.8/110.4 | 153.3 | 148.8 | 118.5 | 119.9 | | | | | | |
| | 133.1/138.1 | 112.8/110.4 | 153.3 | 141.5 | - | - | | | | | | |
| | - | - | - | 150.6 | 116.7 | 120.3 | | | | | | |
| | 131.5 | - | - | - | - | 123.8 | | | | | | |
| | 131.0/133.5 | - | - | - | - | 120.0 | | | | | | |
| | 133.0 | - | - | - | - | 118.0 | | | | | | |
| | 134.3/136.5 | - | - | - | - | 122.6 | | | | | | |

* Some G lignin C3 and C4 have chemical shift values overlapping with S3/5, and these sites are indicated as G'3/4 in the figures.

**Supplementary Table 5. Integration areas for NMR spectral analysis.** $F_2$ = direct $^{13}$C dimension; $F_1$ = indirect $^{13}$C dimension. S/G: deprotonated lignin aromatic carbons, i.e., S(1, 3-5) and G(1,3-4); Carbo: carbohydrate sites including i/s(1-6), Xn(1-5) and GalA(1-5); OMe: lignin methoxy carbons; AcMe: acetyl methyl carbons; CO: carbonyl carbons, including those from acetyl and C6 of GalA. The spectral regions (A1-A4 and B1-B8) are shown in Supplementary Fig. 5.

| Spectra | Integral Name | Integral Region ($F_2$) (ppm) | Integral Region ($F_1$) (ppm) | $^{13}$C sites/<br>$^{13}$C pairs | Integration Analysis | Analysis Description |
|---|---|---|---|---|---|---|
| **1D CP<br>1D DP<br>1D cross-section of gated PDSD** | A1 | 190.0 – 0.0 | - | All $^{13}$C | A3/A1 | 1D CP analysis, rigid lignin content in the whole cell |
| | A2 | 106.0 – 59.0 | - | Carbohydrate $^{13}$C<br>i/s(1-6)<br>Xn(1-5)<br>GalA(1-5) | | |
| | A3 | 156.0 – 109.0 | - | Lignin $^{13}$C*<br>S(1, 3-5)<br>G(1-6) | $\frac{(A4/A1)\ \text{in DP}}{(A4/A3)\ \text{in CP}}$ | 1D DP analysis, quantitative lignin content in the whole cell |
| | A4 | 156.0 – 140.0 | - | S(3/5)<br>G(3/4) | A2/A1 | 1D cross-section of 2D gated PDSD analysis, specific carbon site to carbohydrate transfer |
| **2D Gated PDSD** | B1 | 180.0 – 15.0 | 124.0 – 160.0 | All $^{13}$C propagated from S/G | (B3+B4+B5+B6+B7+B8)/(B1+B2)<br>(B3+B4)/(B1+B2)<br>(B5+B6+B7+B8)/(B1+B2) | (S/G+OMe) – (Carbo+CO+AcMe)<br>(S/G+OMe) – Carbo<br>(S/G+OMe) – (CO+ AcMe) |
| | B2 | 180.0 – 15.0 | 52.0 – 58.5 | All $^{13}$C propagated from OMe | | |
| | B3 | 106.5 – 59.0 | 124.0 – 160.0 | S/G – Carbohydrate | B3/B1<br>(B5+B6)/B1<br>B5/B1<br>B6/B1 | S/G – Carbo<br>S/G – (CO+ AcMe)<br>S/G – CO<br>S/G – AcMe |
| | B4 | 106.5 – 59.0 | 52.0 – 58.5 | OMe – Carbohydrate | | |
| | B5 | 177.0 – 168.0 | 124.0 – 160.0 | S/G – CO | | |
| | B6 | 24.0 – 17.0 | 124.0 – 160.0 | S/G – AcMe | B4/B2<br>(B7+B8)/B2<br>B7/B2<br>B8/B2 | OMe - Carbo<br>OMe – (CO+ AcMe)<br>OMe – CO<br>OMe – AcMe |
| | B7 | 177.0 – 168.0 | 52.0 – 58.5 | OMe – CO | | |
| | B8 | 24.0 – 17.0 | 52.0 – 58.5 | OMe – AcMe | | |
| **2D Gated DARR** | A1 | 106.0 – 103.5 | 172.0 – 174.0 | AcCO – $Xn^{2f}$(1) | A1/(A1+A2+A3) | molar distribution of acetyl groups in $Xn^{2f}$ |
| | A2 | 103.5 – 101.5 | 172.0 – 174.0 | AcCO – $Xn^{3f}$(1) | A2/(A1+A2+A3) | molar distribution of acetyl groups in $Xn^{3f}$ |
| | A3 | 101.5 – 99.5 | 172.0 – 174.0 | AcCO – GalA(1) | A3/(A1+A2+A3) | molar distribution of acetyl groups in GalA |

*Aromatic residue sidechain and fatty acid chain carbon sites in the A3 region of 1D are at 136, 130 and 128.5 ppm and only show up in the 1D DP
